# Extracellular vesicles drive cross-genus mycovirus transmission and suppress two fungal diseases

**DOI:** 10.64898/2026.08.03.742386

**Authors:** Jianing Wu, Jialing Xu, Shutong Yang, Yang Lin, Huang Huang, Jiatao Xie, Jiasen Cheng, Tao Chen, Bo Li, Xiao Yu, Xueliang Lv, Yanping Fu, Xueqiong Xiao, Qing Cai, Daohong Jiang

**Author notes:** Lead contact and correspondence: Yang Lin, Daohong Jiang. These authors contributed equally to this work (Jianing Wu, Jialing Xu, Shutong Yang).

## Abstract

Mycoviruses are typically transmitted vertically through fungal reproduction or horizontally via hyphal anastomosis, but identical viruses in phylogenetically divergent fungi hint at unknown inter-species transmission mechanisms. Here, we identify fungal extracellular vesicles (EVs) as mediators of cross-genus mycovirus transmission. Using the hypovirulent *Botrytis cinerea* strain IBc-374 (harboring 16 mycoviruses) as a donor, we show that up to 13 mycoviruses are horizontally transmitted to *Sclerotinia sclerotiorum* during dual culture or plant co-inoculation, even though these two fungi belong to different genera and are generally considered incapable of hyphal anastomosis. Electron microscopy reveals abundant vesicle structures in IBc-374 hyphae, and nanoparticle tracking analysis showed that the strain secretes >11-fold more EVs than a virus-free strain. RT-PCR detects genomic RNAs of 7 mycoviruses in purified EVs, and the full-length viral genome in EVs was further validated with BcHV5 as example by fluorescence in situ hybridization and RT-PCR. Incubation of protoplast-derived germlings of *S. sclerotiorum* or *B. cinerea* with IBc-374 EVs leads to infection by 4–6 donor mycoviruses, demonstrating EV-mediated cross-genus transmission. Injection of mycovirus-carrying EVs into tobacco leaves followed by fungal inoculation also transmits two hypoviruses to both species. Application of IBc-374 hyphal fragment suspension significantly reduces lesion sizes caused by both pathogens on plants, and rescued two pathogens carrying multiple mycoviruses exhibit hypovirulence and impaired growth. Our findings reveal EVs as a cell-free vector that bypasses vegetative incompatibility, providing a mechanistic basis for cross-species viral spread and opening avenues for EV-based virus cocktails to control multiple fungal diseases.

**IN BRIEF:** Wu et al. discover that fungal extracellular vesicles (EVs) can package and transmit multiple mycoviruses across genera from *Botrytis cinerea* to *Sclerotinia sclerotiorum*. EV-mediated delivery overcomes vegetative incompatibility barriers and reduces disease caused by both fungal pathogens, offering a cell-free strategy for mycovirus-based biological control.

## INTRODUCTION

Mycoviruses are ubiquitous in fungi and oomycetes, and certain hypovirulence-associated mycoviruses offer promising agents for biological control of plant diseases^1–3^. The successful application of Cryphonectria hypovirus 1 (CHV1) against chestnut blight and Sclerotinia sclerotiorum hypovirulence-associated DNA virus 1 (SsHADV1) against white mould exemplify this potential^4–6^. However, the widespread use of mycoviruses is constrained by their often inefficient horizontal transmission with rare exception^7^, which largely depends on hyphal anastomosis. Vegetative incompatibility systems in many fungi pose a major barrier to hyphal fusion^8^, thereby restricting mycovirus spread^9^. Strategies to overcome this barrier include engineering “super-donor” strains^10^ or using chemicals to attenuate incompatibility responses^11,12^. A transformative alternative would be a host-cell-independent delivery mechanism that bypasses vegetative incompatibility altogether.

Extracellular vesicles (EVs) are membrane-derived nanostructures released by virtually all cell types and are recognized as key mediators of intercellular communication^13–16^. EVs transport diverse cargoes, such as proteins, lipids, and RNAs, and influence recipient cell behaviour^17–19^. Importantly, several animal and plant viruses are packaged into EVs, which can facilitate viral dissemination^20–26^. For fungi, membrane-associated particles containing CHV1 genomic RNA and RNA polymerase activity were isolated from *C. parasitica* decades ago^27,28^, but whether mycoviruses are actively loaded into fungal EVs and whether such EVs mediate mycovirus transmission has remained unexplored^29^.

Intriguingly, accumulating evidence suggests that mycoviruses can undergo cross-species transmission, including between distantly related fungi. For example, high sequence similarity has been observed between Botrytis porri RNA virus 1 and Sclerotinia sclerotiorum botybirnavirus 3, and shared viral sequences occur between *Botrytis cinerea* and *Sclerotinia sclerotiorum*, two cosmoplitan plant fungal pathogens that belong to the same family^30–33^. Cross-class and even cross-phylum transmissions have also been reported^34,35^, indicating that mycovirus host ranges may be less constrained than previously thought. The mechanisms underlying these transmission events, however, remain largely unknown. Given the ability of EVs to be taken up by diverse cell types^36–39^, we hypothesized that EVs might represent a novel vehicle for cross-genus mycovirus transmission.

In this study, we investigated the hypovirulent *B. cinerea* strain IBc-374, which is co-infected by 16 mycoviruses^40^. We show that multiple mycoviruses can be transmitted from IBc-374 to strain Ep-1PNA367^R^ of *S. sclerotiorum*, an intergeneric fungus of *B.cinerea*. Crucially, we discover that IBc-374 secretes an elevated amount of EVs, and that several mycovirus genomes are encapsulated within these EVs. Purified EVs can be internalized by protoplast-derived germlings of both *S. sclerotiorum* and *B. cinerea*, leading to productive viral infection. Furthermore, EV-mediated transmission occurs in planta and confers hypovirulence to recipient *S. sclerotiorum* and *B. cinerea*. Finally, we demonstrate that hyphal suspensions of IBc-374 reduce disease caused by both pathogens, partly through EV-mediated virus spread and partly through the induction of plant defence responses. Our findings reveal EVs as an unrecognized vehicle for cross-genus mycovirus transmission and open new avenues for mycovirus-based biological control of fungal diseases.

## RESULTS

### Sixteen Mycoviruses Co-infect the Hypovirulent *B. cinerea* Strain IBc-374

Strain IBc-374 exhibited altered colony morphology compared to the virulent strain B05.10, displaying regular colony margins and significantly reduced colony size with shorter, denser aerial mycelia (Figure 1A). Quantitatively, the hyphal growth rate of IBc-374 was 7.8 ± 0.23 mm/day, a 46.58% reduction compared to B05.10 (14.6 ± 0.35 mm/day) (Figure 1A; t-test, *P* < 0.001, *n* = 4). Furthermore, IBc-374 was avirulent, failing to induce lesion formation on tobacco leaves at 72 hours post-inoculation (hpi), whereas B05.10 caused robust infection (Figure 1B; t-test, *P* < 0.001, *n* = 5). Transmission electron microscopy revealed extensive organelle disruption in IBc-374 hyphae, while B05.10 hyphae maintained intact cellular structures under identical culture conditions (Figure 1C, D).

**Figure 1.**
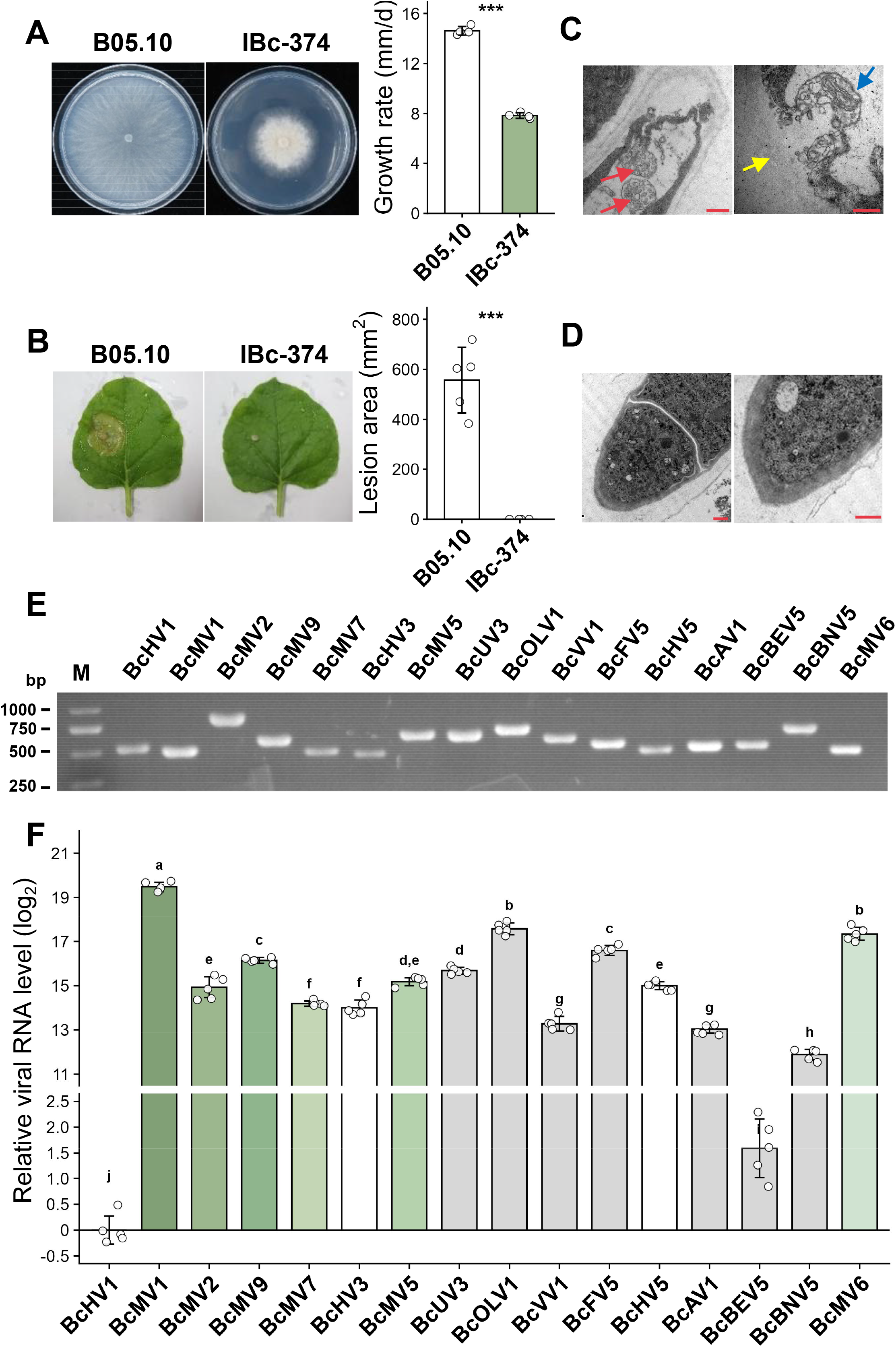
Biological characterization and infected mycoviruses of *B. cinerea* hypovirulent strain IBc-374. **(A)** The colony morphology and growth rate of *B. cinerea*. Images were taken at 3 days post inoculation (dpi) on PDA at 20℃. Growth rate of *B. cinerea* strains was calculated as the increase in colony radius between 1 and 2 dpi. Colony diameters were measured at 1 and 2 dpi on PDA at 20℃. Data are from four independent replicates. Error bars indicate standard deviation (SD). Statistical significance was determined by Student’s t test (\*\*\**P* < 0.001). **(B)** Lesion induced by *B. cinerea* strains on detached tobacco leaves (20℃, 3 dpi). Photographs were captured at 3 dpi. Lesion area were measured and statistically analyzed after incubation at 20℃ for 3 dpi. Data are from five independent replicates. Error bars indicate standard deviation (SD). Statistical significance was determined by Student’s t test (\*\*\**P* < 0.001). **(C)** TEM image of strain IBc-374. Red arrows indicate multivesicular body (MVB)-like structures. The blue arrow indicates numerous intracellular vesicles released from the cell membrane. The yellow arrow indicates the cell wall. Scale bar, 0.5 µm. **(D)** TEM image of strain B05.10. Scale bar, 1 µm. **(E)** Mycoviruses harbored by strain IBc-374. RT-PCR analysis of mycoviruses in strain IBc-374. The presence of viral RNAs was detected using virus-specific primers. **(F)** Relative viral RNA levels in strain IBc-374. Real-time PCR was performed using mycovirus-specific primers. Relative viral RNA levels were calculated using the 2^-ΔΔCt^ method, and log_2_-transformed values are shown. Data represent five independent biological replicates. Error bars indicate standard deviation (SD). Statistical analysis was performed using ANOVA, followed by Tukey’s post-hoc test, with statistical significance set at *P* < 0.01.

Virome sequencing and RT-PCR confirmed the co-infection of IBc-374 by 16 mycoviruses (Figure 1E, Table S1). This viral assemblage comprised six mitoviruses (BcMV1, BcMV2, BcMV5, BcMV6, BcMV7, and BcMV9), one splipalmivirus (BcBNV5), one ourmia-like virus (BcOLV1), one betaendornavirus (BcBEV5), one fusarivirus (BcFV5), three hypoviruses (BcHV1, BcHV3, and BcHV5), one mycoalphavirus (BcAV1), one umbra-like virus (BcUV3), and one victorivirus (BcVV1). Nucleotide sequences for all mycoviruses have been deposited in NCBI GenBank (Table S1). The RNA polymerase of BcBEV5 shared 67.7% amino acid identity and 100% coverage with that of Botrytis cinerea betaendornavirus 1. Phylogenetic analysis classified these mycoviruses into nine established families: *Mitoviridae*, *Splipalmiviridae*, *Botourmiaviridae*, *Hypoviridae*, *Ambiguiviridae*, *Fusariviridae*, *Pseudototiviridae*, *Mycoalphaviridae,* and *Endornaviridae* (Figure S1). These results demonstrate that the hypovirulent *B. cinerea* strain IBc-374 is co-infected by a diverse community of mycoviruses.

The relative abundance of each viral RNA within IBc-374 was determined by RT-qPCR. BcMV1 exhibited the highest RNA level, followed by BcMV6 and BcOLV1. Conversely, BcHV1 displayed the lowest RNA level, followed by BcBEV5 and BcBNV5 (Figure 1F) (ANOVA, Tukey’s post-hoc test, *P* < 0.01, *n* = 5).

### Mycoviruses Fail to Transmit from Strain IBc-374 to the Vegetatively Incompatible Strain B05.10^G^ on PDA

To assess the horizontal transmission potential of mycoviruses from IBc-374, dual-culture experiments on PDA were performed using the vegetatively incompatible strain B05.10^G^ as a recipient (Figure 2A). Following these attempts, 15 subcultures from dual-culture experiments were selected on hygromycin-amended PDA. RT-PCR analysis revealed that none of the 16 mycoviruses were successfully transmitted from IBc-374 to B05.10^G^ in vitro. These negative results were consistently reproduced across multiple independent replicates. Evans blue staining indicated that IBc-374 and B05.10^G^ belong to different vegetative compatibility groups (VCGs) (Figure 2B), suggesting that vegetative incompatibility prevents mycovirus transmission via hyphal anastomosis.

**Figure 2.**
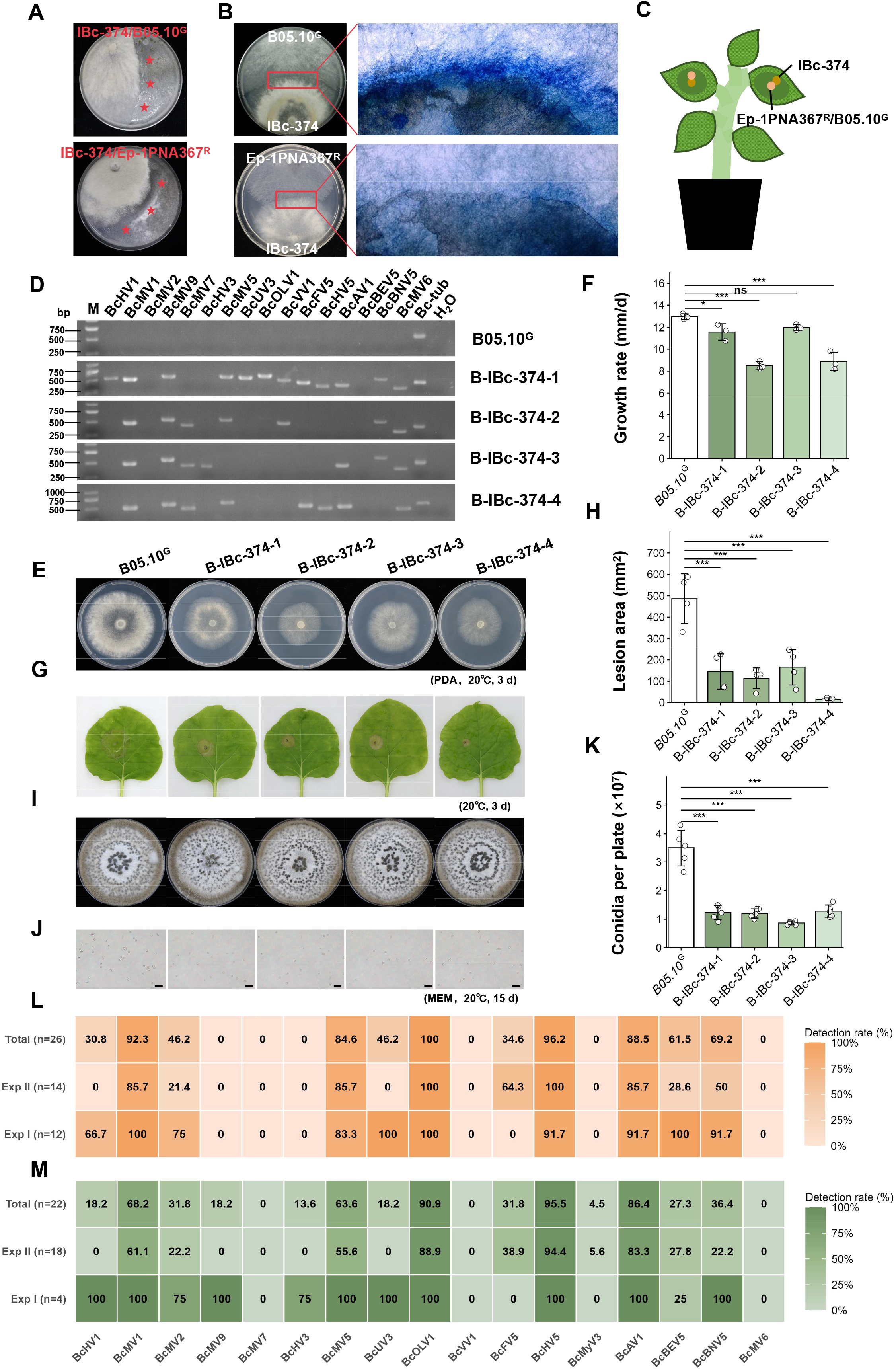
Horizontal transmission of mycoviruses from *B. cinerea* hypovirulent strain IBc-374 to *S. sclerotiorum* strain Ep-1PNA367^R^ or *B. cinerea* strain B05.10^G^. **(A)** Dual culture of strains IBc-374 with B05.10^G^ or Ep-1PNA367^R^ on PDA at 20℃ for 7 days. Red asterisks indicate the sites from which subcultures were isolated. **(B)** Evans blue staining of strains IBc-374 and B05.10^G^ or Ep-1PNA367^R^ after co-cultured on PDA for 20 h at 20℃ to detect programmed cell death at the mycelial interface. Mycelium within the red box areas was stained with 0.5% (w/v) Evans blue solution for 30 min at 20℃. **(C)** Co-inoculation of strain IBc-374 with Ep-1PNA367^R^ or B05.10^G^ on *Nicotiana benthamiana* leaves under 100% relative humidity in a greenhouse. Recipient strains were rescued from lesions and cultured on PDA supplemented with 50 μg/μL hygromycin B (or neomycin) and 100 μg/μL cephalosporin. **(D-I)** Horizontal transmission of IBc-374 mycoviruses to *B. cinerea* via sprayed mycelial suspension on *N. benthamiana*. B05.10^G^ were inoculated onto *N. benthamiana* leaves after pretreatment with IBc-374 hyphal suspension. Plants were maintained at 20℃ under high humidity (100%). B05.10^G^ subcultures were rescued from lesions of inoculated *N. benthamiana* leaves. **(D)** Mycoirus profiles of subcultures examined by RT-PCR. *Tubulin* of *B. cinerea* was used as a positive control for PCR amplification, and H_2_O was used as a negative control for the PCR template. **(E)** Colony morphology of four representative subcultures. Images were taken at 72 hpi on PDA at 20℃. **(F)** Average mycelial growth rate of four subcultures. Colony diameters were measured at 24 and 48 hpi. Growth rate of each strain was calculated. **(G, H)** Pathogenicity of the four subcultures on detached *N. benthamiana* leaves. **(G)** Representative lesions at 48 hpi (20℃). **(H)** Lesion area of four subcultures on detached rapeseed leaves (20℃, 48 hpi). **(I-K)** Conidiation of four subcultures on MEM plates after 15 days at 20℃. **(I)** Sporulation on MEM plates. **(J)** Conidial morphology. **(K)** Spore yield per plate. Strains designated B-IBc-374-1, B-IBc-374-2, B-IBc-374-3, and B-IBc-374-4 were *B. cinerea* isolates recovered from plants that had been sprayed with the IBc-374 mycelial suspension. Each strain had four replicates and the experiment was repeated twice. Error bars indicate standard deviation (SD). The data were analyzed using Dunnett’s test (\*\**P* < 0.01; \*\*\**P* < 0.001; ns not significant). **(L)** Heatmap illustrating the efficiency of mycovirus transmission from *B. cinerea* to *S. sclerotiorum* on PDA. Recipient *S. sclerotiorum* strains were isolated from dual cultures, and virus acquisition was assessed by RT-PCR using virus-specific primers. Two independent experiments were performed, yielding 12 and 14 derivative strains, respectively. Each row represents an individual independent experiment, and each column represents a specific mycovirus. The color intensity reflects the transmission efficiency, calculated as the proportion of recipient strains that acquired each virus within a given experiment. **(M)** Heatmap illustrating the efficiency of mycovirus transmission from *B. cinerea* to *S. sclerotiorum* on *N. benthamiana* leaves. Recipient strains were isolated from co-inoculated leaves, and virus acquisition was assessed by RT-PCR. Two independent experiments were performed, yielding 4 and 18 derivative strains, respectively. Each row represents an individual independent experiment, and each column represents a specific mycovirus. The color intensity reflects the transmission efficiency, calculated as the proportion of recipient strains that acquired each virus within a given experiment.

### Mycoviruses are Transmitted from Strain IBc-374 to *B. cinerea* Strain B05.10^G^ in planta

To probe if viruses can transmit between two vegetative incompatible strains, the hyphal fragments of IBC-374 were sprayed on *Nicotiana benthamiana* leaves, and then B05.10^G^ was inculated on the same leaves (Figure 2C), after lesion formed, the lesion margins of *N. benthamiana* were taken for reisolating B05.10^G^ using neomycin ammending PDA plates, four derivative strains of *B. cinerea* B05.10^G^ were recovered. Unlike on PDA no virus can transmit from IBC-374 to *B. cinerea* B05.10G, RT-PCR analysis revealed that these derivatives acquired up to 14 mycoviruses, including BcHV1, BcMV1, BcMV9, BcMV7, BcHV3, BcMV5, BcUV3, BcOLV1, BcVV1, BcFV5, BcHV5, BcAV1, BcBNV5, and BcMV6 (Figure 2D). Phenotypic characterization further demonstrated that these virus-carrying *B. cinerea* strains exhibited varying degrees of debilitation, including reduced mycelial growth, attenuated virulence, and decreased sporulation capacity (Figure 2E-K). In virulence assays on tobacco leaves, all four virus-infected strains showed significantly reduced pathogenicity compared with the control strain B05.10^G^ (mean lesion area of 486.12 mm^2^). The mean lesion areas of B-IBc-374-1, B-IBc-374-2, B-IBc-374-3, and B-IBc-374-4 were 145.40, 113.14, 165.63, and 15.28 mm^2^, corresponding to reductions of 70.09%, 76.73%, 65.93%, and 96.85%, respectively (Figure 2G, H). Notably, B-IBc-374-4 exhibited the most severe attenuation, with its lesion area almost completely eliminated. Similarly, sporulation capacity was also dramatically impaired, as all virus-infected strains produced significantly fewer spores than the control (Dunnett’s test, *P* < 0.001 for all strains, Figure 2I-K). The reduction ranged from 63.27% (B-IBc-374-4) to 75.29% (B-IBc-374-3). Collectively, these findings indicate that strain IBc-374 is capable of transmitting mycoviruses to *B. cinerea* in planta.

### Mycoviruses are Transmitted from Strain IBc-374 to *S. sclerotiorum* Strain Ep-1PNA367^R^ Both on PDA and in Planta

We found that mycoviruses were successfully transmitted from IBc-374 to the intergeneric *S. sclerotiorum* strain Ep-1PNA367^R^ in both dual-culture experiments on PDA and co-inoculation assays on plant leaves (Figure 2). In vitro dual-culture on PDA yielded 26 recovered Ep-1PNA367^R^ subcultures, and RT-PCR analysis confirmed the transmission of 11 mycoviruses: BcHV1, BcMV1/2/5, BcUV3, BcOLV1, BcFV5, BcHV5, BcAV1, BcBEV5, and BcBNV5 (Figure 2L). The remaining five viruses (BcMV6/7/9, BcHV3, and BcVV1) were not detected in the *S. sclerotiorum* subcultures. In planta co-inoculation experiments on *N. benthamiana* (Figure 2C) yielded 22 viable Ep-1PNA367^R^ subcultures from lesion margins, with cross-genus transmission observed for 13 mycoviruses, including all 11 mycoviruses transmitted on PDA plus two additional mycoviruses (BcMV9 and BcHV3) (Figure 2M). Notably, *S. sclerotiorum* and *B. cinerea* belong to phylogenetically distinct genera and are incapable of typical hyphal anastomosis. Although cell death occurred at the hyphal interaction zones of the two fungal species (Figure 2B), it was not due to vegetative incompatibility reaction. Moreover, the extent of cell death at the contact zone of the two colonies in the IBc-374/Ep-1PNA367^R^ pairing was significantly lower than that observed in the IBc-374/B05.10^G^ pairing, which was initiated by vegetative incompatibility reaction, as evidenced by the differential staining intensity at the hyphal contact zones (Figure 2B). These results suggest that an anastomosis-independent mechanism facilitates intergeneric viral transmission.

### Characteristics of Extracellular Vesicles (EVs) Secreted by *B. cinerea* IBc-374

To investigate the potential role of EVs in the intergeneric spread of mycoviruses, we first characterized the EVs secreted by the mycovirus-infected *B. cinerea* strain IBc-374, using the virus-free strain B05.10 as a control. Transmission electron microscopy (TEM) of purified nanoparticles from culture filtrates of both strains revealed vesicles with the characteristic cup-shaped morphology of EVs (Figure 3A). EVs were purified from 300 mL of culture filtrate and resuspended in 1000 μL of 1 × PBS. Nanoparticle tracking analysis (NTA) showed that strain IBc-374 produced EVs at a concentration of (7.63 ± 0.15) × 10^11^ particles/mL, with a median particle size of 126.2 ± 2.2 nm. In contrast, the control strain B05.10 produced EVs at a significantly lower concentration of (6.77 ± 0.06) × 10^10^ particles/mL, with a larger median particle size of 148.2 ± 1.6 nm (Figure 3B, C). This represents an 11.3-fold increase in EV production by the mycovirus-infected hypovirulent strain IBc-374 compared to the uninfected virulent strain B05.10 (t-test, *P* < 0.001, *n* = 3). These findings indicate that IBc-374 produces significantly more EVs with smaller sizes than B05.10. (t-test, *P* < 0.001, *n* = 3). These combined ultrastructural and biophysical characteristics are consistent with the established hallmarks of exosomes^41^, suggesting that mycovirus infection in strain IBc-374 is associated with enhanced production of EVs.

**Figure 3.**
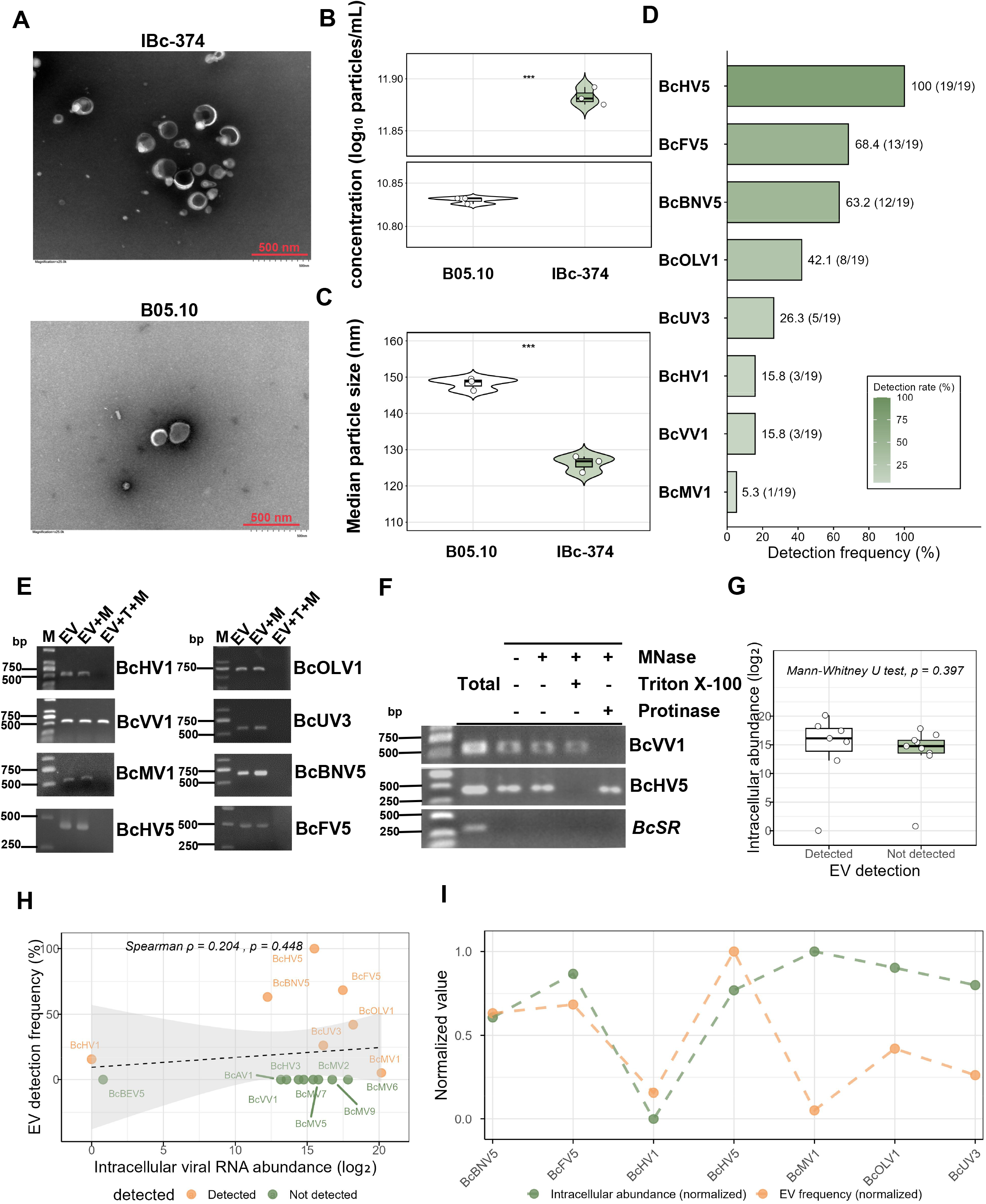
Mycoviruses are selectively packaged within EVs of *B. cinerea* hypovirulent strain IBc-374. **(A)** Transmission electron microscope (TEM) image of EVs purified from the culture filtrate of stationary-cultured *B. cinerea* strains. Scale bar = 1 µm. **(B-D)** Characterization of EVs purified from *B. cinerea* strains IBc-374 and B05.10. EVs were purified from stationary-cultured *B. cinerea* strains. Three replicates were performed per sample. EVs from *B. cinerea* strains were characterized by nanoparticle tracking analysis (NTA). **(B)** Violin plot of EV concentration from *B. cinerea* strains. Boxplots within the violins indicate the median and interquartile range, and individual data points are overlaid as black circles. Data are from three independent replicates. Statistical significance was determined by Student’s t test (\*\*\**P* < 0.001). **(C)** Violin plot of median particle size of EVs from *B. cinerea* strains. Boxplots within the violins indicate the median and interquartile range, and individual data points are overlaid as black circles. Data are from three independent replicates. Statistical significance was determined by Student’s t test (***P < 0.001). **(D)** Detection frequency of mycoviruses in EVs of strain IBc-374. EVs were independently isolated 19 times. The presence of mycoviruses in EV preparations was detected by RT-PCR using virus-specific primers, and the detection frequency was calculated for each virus. **(E)** Encapsulation of mycoviruses within EVs secreted by strain IBc-374. Nuclease protection assays were performed to determine whether viral RNAs were protected by EV membranes. After each treatment, viral RNAs were detected by RT-PCR. “EV” represents extracellular vesicles without any treatment, “EV + M” represents extracellular vesicles treated with MNase, and “EV + T + M” represents extracellular vesicles treated with Triton X-100 and MNase. **(F)** BcVV1 exists as free viral particles. Enzymatic treatments were used to determine whether BcVV1 viral particles are encapsulated within EVs. EVs were divided into 250 µL aliquots and subjected to four conditions: (i) untreated control; (ii) MNase alone; (iii) Triton X-100 (30 min) followed by MNase; (iv) Proteinase K (60 min at 37 ℃) followed by PMSF inactivation and then MNase. After treatments, RNA was extracted and analyzed by RT-PCR using BcVV1-specific primers. *B. cinerea* transcriptional regulator gene *BcS*R was used as a negative control for cytoplasmic contamination, and mycovirus BcHV5 (which lacks viral particles) was used as a positive control. **(G)** Abundance comparison between EV-detected and non-detected viruses. Boxplot comparing log_2_-transformed intracellular viral RNA abundance between EV-detected (n = 7, red) and non-detected (n = 10, blue) viruses. Boxes represent the interquartile range (IQR), horizontal lines indicate medians, and whiskers extend to the full data range. Individual data points are overlaid as black jittered points. No significant difference in abundance was observed between the two groups (Mann-Whitney U test, *P* = 0.397). **(H)** Intracellular abundance does not correlate with EV packaging frequency. Scatter plot showing the relationship between log_2_-transformed intracellular viral RNA abundance and EV detection frequency (%) for 16 mycoviruses. Orange circles: viruses detected in EVs (n = 7). Green circles: viruses not detected in EVs (n = 9). The dashed line and gray shaded area represent the linear regression line and 95% confidence interval, respectively. Spearman rank correlation analysis revealed no significant monotonic relationship (ρ = 0.204, *P* = 0.448). **(I)** Normalized contrast of abundance and EV frequency among detected mycoviruses. For the seven EV-detected mycoviruses, intracellular abundance (green) and EV detection frequency (orange) were normalized to a 0–1 scale to compare their relative patterns. Points are connected by dashed lines for each virus. No consistent direct or inverse relationship is observed.

### Mycoviruses are Selectively Packaged within EVs of Strain IBc-374

To determine whether mycoviruses are packaged within EVs, we isolated EVs from the culture filtrate of IBc-374 and performed RT-PCR to detect the presence of viral RNAs. We successfully detected eight mycoviruses (BcHV1, BcUV3, BcBNV5, BcHV5, BcOLV1, BcFV5, BcMV1, BcVV1) within the EVs samples. To confirm that the viral RNAs were encapsulated within the EVs and not simply adhering to the surface, we performed nuclease protection assays. Viral RNAs were resistant to digestion by micrococcal nuclease (MNase) alone, but became susceptible to MNase digestion following disruption of the EV membrane with Triton X-100 (Figure 3E). This confirms that the viral RNAs are protected within the EV lumen. Notably, while the nucleic acid of mycovirus BcVV1 was detectable in the EV suspension, it was insensitive to perforator treatment. To determine whether the viral particles of BcVV1 are encapsulated within EVs, we further designed treatment assays combining nuclease, perforator, and proteinase K. The results showed that under co-treatment with nuclease and proteinase K, the nucleic acid of BcVV1 was completely degraded even in the absence of perforator, indicating that the viral particles of BcVV1 are not encapsulated within EVs (Figure 3F). When treated simultaneously with perforator and nuclease, the nucleic acid of the control BcHV5 could not be detected regardless of whether proteinase K was applied. Furthermore, no nucleic acid of the highly expressed transcriptional regulator gene, *B. cinerea* Sterol Regulator (BcSR, BCIN_02g03500)^42^ was detected in the RNA extracted from EVs in this assay, indicating that the extracted EVs were not contaminated with cytoplasmic RNA. The detection frequencies of these mycoviruses varied significantly across 19 independent EV preparations, with BcHV5 exhibiting the highest detection rate (100%) and BcMV1 the lowest (5.3%) (Figure 3D).

To determine whether the intracellular abundance of mycoviruses influences their encapsulation into EVs, we compared the intracellular abundance distributions of EV-detected and non-detected viruses. The boxplot (Figure 3G) showed no significant difference between the two groups (median log₂ abundance: 16.12 vs 14.76, respectively; Mann-Whitney U test, *P* = 0.397). The abundance ranges overlapped extensively (detected: 0.003–20.16; non-detected: 0.8–17.84), and individual data points were intermingled between groups. This result demonstrates that intracellular abundance does not predict whether a virus is packaged into EVs. Scatter plot analysis revealed extensive overlap in intracellular abundance between EV-detected (orange circles, *n* = 7) and non-detected viruses (green circles, *n* = 10) (Figure 3H). Spearman rank correlation showed no significant monotonic relationship between intracellular abundance and EV detection frequency across all 16 mycoviruses (ρ = 0.204, *P* = 0.448). Notably, BcMV1, which exhibited the highest abundance (log_2_ fold change = 19.53), was detected at a low frequency (5.3%), whereas BcHV5, with moderate - high abundance (log_2_ fold change = 15.01), was detected in every EV sample (100%). Conversely, BcHV1, which had the lowest abundance (set as reference; log_2_ fold change = 0.00), showed an intermediate detection frequency (15.8%). These findings indicate that EV packaging is not a passive, abundance-driven process. Among the seven EV-detected viruses, four — BcHV5, BcFV5, BcBNV5, and BcHV1 — exhibited a positive correspondence between their intracellular abundance and EV detection frequency, whereas the remaining three (BcMV1, BcOLV1, and BcUV3) deviated from this trend (Figure 3I). Specifically, BcHV5, which possessed the highest abundance among this subset (log_2_ fold change = 15.01), showed the highest EV detection frequency (100%). BcFV5 (log_2_ fold change = 17.48) and BcBNV5 (log_2_ fold change = 11.93) displayed proportionally high (68.4%) and moderate (63.2%) frequencies, respectively. At the lower end, BcHV1, with extremely low abundance (log_2_ fold change = 0.00), showed the lowest detection frequency (15.8%). In contrast, the remaining three viruses—BcMV1, BcOLV1, and BcUV3—did not follow this pattern. Notably, BcMV1, which had the highest overall abundance (log_2_ fold change = 19.53), showed the lowest EV frequency (5.3%), while BcOLV1 and BcUV3 displayed intermediate frequencies (42.1% and 26.3%, respectively) that were lower than expected based on their abundance ranks.

To further confirm the encapsulation of mycoviruses within EVs, we performed fluorescence in situ hybridization (FISH) to detect the viral RNA of BcHV5 which has the largest genome size among detected viruses in EVs. Co-staining with the lipophilic dye FM4-64 revealed precise co-localization between EV membranes (red fluorescence) and viral RNA (green fluorescence pseudo-colored) via confocal microscopy (Figure 4A). Complete genome sequencing of EV-packaged BcHV5, using segment-specific primers, showed identical amplification profiles to intracellular viral RNA (Figure 4B), demonstrating intact genomic packaging. Northern blot analysis, using a BcHV5-specific probe, revealed a band of the same size as that detected in the total RNA of strain IBc-374 in the nucleic acids extracted from EVs, further confirming that the EVs secreted from strain IBc-374 carry the complete nucleic acid sequence of the mycovirus BcHV5 (Figure 4C).

**Figure 4.**
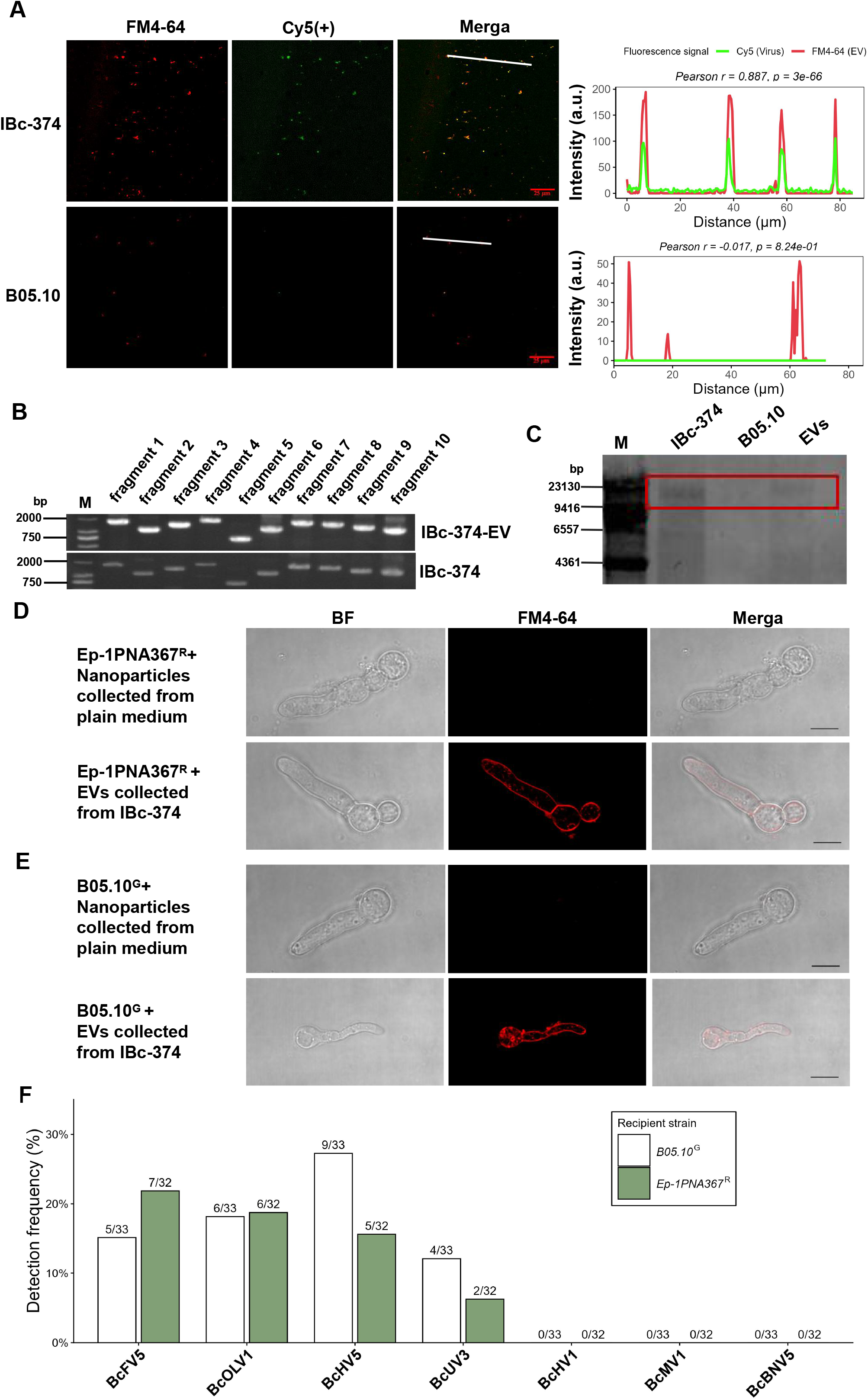
EVs containing mycoviruses can be internalized by fungi and mediate virus transmission in vitro. **(A)** Co-localization of BcHV5 with EVs. FM4-64 labels EVs (red); a Cy5-conjugated BcHV5-specific probe appears green (pseudo-colored). To resolve spectral overlap, FM4-64 was excited at 561 nm (offset -10) with emission at 570–616 nm, and Cy5 at 638 nm (offset 0) with emission at 633–738 nm. Negative offset and spectral unmixing were applied to minimize crosstalk. White lines indicate transects for fluorescence intensity measurements; red and green lines represent FM4-64 and Cy5 fluorescence intensities, respectively. Scale bar, 25 µm. Three replicates were performed. **(B, C)** EVs of strain IBc-374 carry the full-length BcHV5 nucleic acid sequence. **(B)** Segmented RT-PCR amplification of the full-length BcHV5 nucleic acid from EV RNA. **(C)** Northern blot detection of BcHV5 in EVs. A DIG-labeled probe prepared with virus-specific primers was hybridized to EV RNA. Signals were detected using NBT/BCIP substrate (purple-blue precipitates). Total RNA from virus-carrying strain IBc-374 served as a positive control; total RNA from virus-free strain B05.10 served as a negative control. **(D, E)** Uptake of EVs by recipient fungi in vitro. **(D)** Uptake of EVs by *S. sclerotiorum* strain Ep-1PNA367^R^. **(E)** Uptake of EVs by *B. cinerea* strain B05.10^G^. FM4-64-labeled EVs were co-incubated with protoplast-derived germlings for 4 h at 20℃, washed, and then imaged by confocal microscopy. PBS stained with FM4-64 served as a negative control. Scale bar, 10 µm. Three replicates were performed per strain. **(F)** Efficiency of viruses acquisition by recipient fungi in vitro. Viral nucleic acids in recipient strains were detected by RT-PCR. A total of 32 *S. sclerotiorum* and 33 *B. cinerea* recipient strains were tested for virus acquisition.

These findings demonstrate that strain IBc-374 actively packages multiple mycoviruses into secreted EVs through a non-random process, as evidenced by the selective packaging efficiency, complete genome preservation, and the inverse correlation between intracellular viral RNA abundance and EV packaging frequency. This suggests a specific mechanism for viral RNA sorting into EVs.

### EVs Containing Mycovirus Can Be Internalized by Fungi

To investigate the potential for EV-mediated mycovirus transmission across fungal species, we developed a uptake assay using fluorescently labeled EVs. EVs were purified from *B. cinerea* strain IBc-374 cultures and labeled with the lipophilic dye FM4-64. These labeled EVs were then incubated with protoplast-derived young hyphae of *S. sclerotiorum* strain Ep-1PNA367^R^ or *B. cinerea* strain B05.10^G^. Medium-derived nanoparticles served as a negative control for non-specific uptake. Confocal microscopy revealed distinct punctate and diffuse fluorescence signals within the cytoplasm of both recipient species at 4 hours post-incubation (hpi), indicative of EV internalization. In contrast, control hyphae exhibited minimal fluorescence signal (Figure 4D, E). These observations suggest active internalization of EVs by both *S. sclerotiorum* and *B. cinerea*.

To assess whether internalized EVs resulted in mycovirus transmission, incubated protoplast-derived hyphae were stringently washed and cultured on PDA. Viral transmission was evaluated in 32 *S. sclerotiorum* (Ep-1PNA367^R^) and 33 *B. cinerea* (B05.10^G^) isolates using RT-PCR. The results demonstrated that both fungal species acquired multiple mycoviruses present in IBc-374-derived EVs. In *S. sclerotiorum* recipients, the infection rates were: BcUV3 (6.3%), BcOLV1 (18.8%), BcFV5 (21.9%), and BcHV5 (15.6%). In *B. cinerea* recipients, the infection rates were: BcUV3 (12.1%), BcOLV1 (18.2%), BcFV5 (15.2%), and BcHV5 (27.3%). To determine whether recipient species influences EV-mediated virus transmission, we compared transmission rates of four transmissible viruses (BcUV3, BcOLV1, BcFV5, and BcHV5) between *S. sclerotiorum* and *B. cinerea* recipients. Fisher’s exact tests revealed no significant differences for any individual virus (all *P* > 0.05). Logistic regression analysis showed no significant effects of virus identity (all *P* > 0.23) or recipient species (*P* = 0.49) on transmission success. A likelihood ratio test further confirmed that virus-recipient interactions did not significantly improve model fit (*P* = 0.5345). Collectively, these results indicate that the four transmissible viruses do not exhibit recipient species-specific transmission patterns. Notably, BcHV1, BcMV1, and BcBNV5 were not detected in any of the recipient strains (Figure 4F).

### EVs-Mediated Cross-Genus Transmission Between *S. sclerotiorum* and *B. cinerea* on Plant

To investigate EV-mediated mycovirus transmission in planta, we injected *B. cinerea* IBc-374-derived EV suspensions into *N benthamiana* leaves, followed by inoculation with *S. sclerotiorum* (Ep-1PNA367^R^) and *B. cinerea* (B05.10^G^) (Figure 5A). At 72 hpi, the inoculated strains were re-isolated from the lesion margins and subjected to mycovirus detection after subculturing. RT-PCR analysis of the purified EVs from IBc-374 confirmed the presence of BcHV1, BcFV5 and BcHV5 (Figure 5B). While BcVV1 was detected in the EV suspension, nuclease protection assays demonstrated that it exists as free viral particles rather than being encapsulated within EVs (Figure 3E).

**Figure 5.**
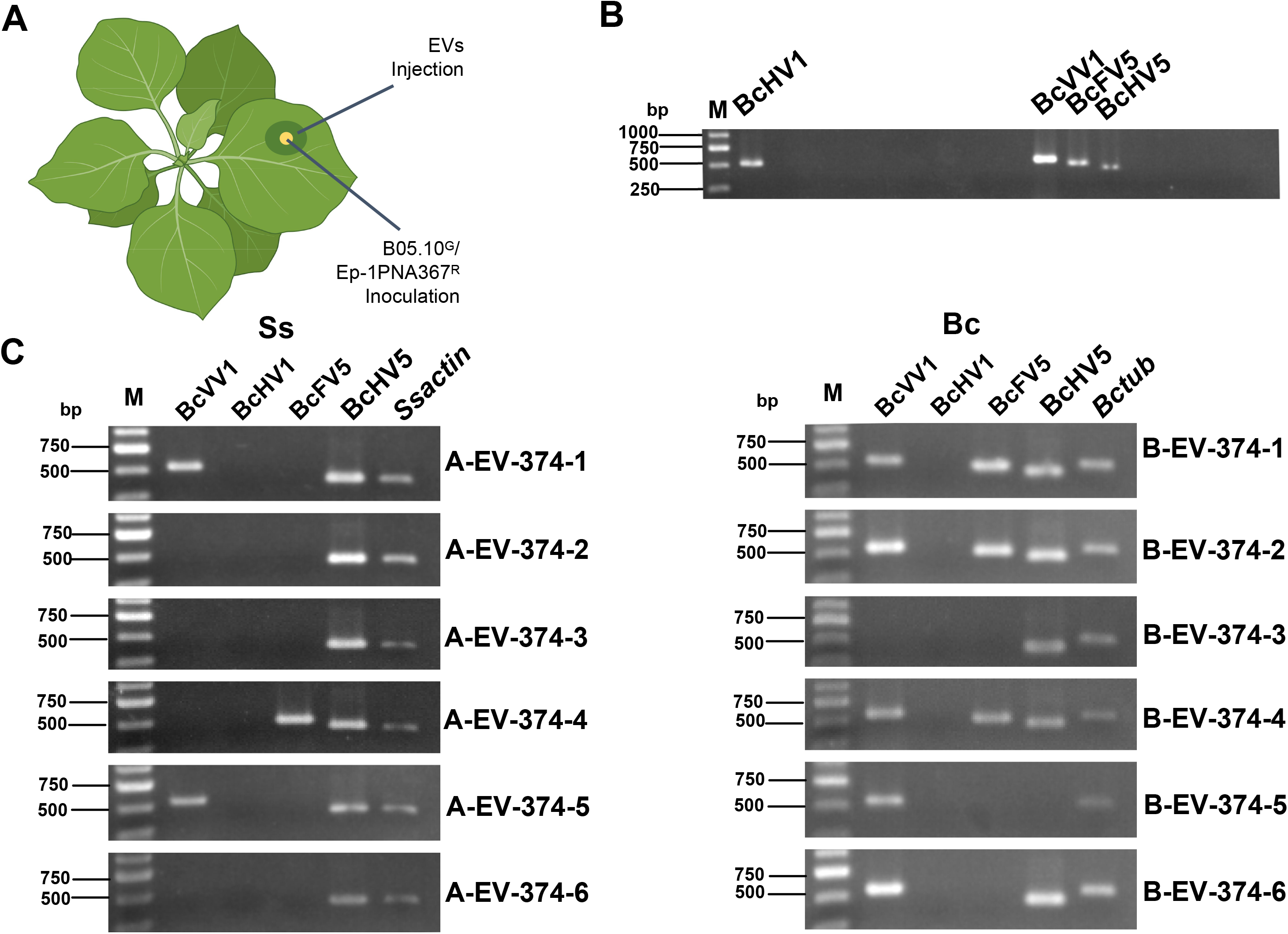
EVs containing mycoviruses mediate virus transmission to recipient fungi in planta. **(A)** Schematic diagram of the experimental workflow. EV suspensions were injected into tobacco leaves, followed by inoculation with *S. sclerotiorum* or *B. cinerea*. At 72 hpi, two recipient strains were re-isolated from lesion margins and analyzed after subculturing. **(B)** Viral profile of the injected EV suspensionas determined by RT-PCR. **(C)** RT-PCR detection of mycoviruses in re-isolated recipient strains using virus-specific primers. Bc: re-isolated *B. cinerea*; Ss: re-isolated *S. sclerotiorum.* Three replicates were performed per treatment.

Six derivative strains each of *S. sclerotiorum* and *B. cinerea* were isolated from the lesion margins. After two rounds of subculturing, the mycovirus profiles of these derivative strains were determined. All six *S. sclerotiorum* derivatives acquired BcHV5, and one strain (A-EV-374-4) also acquired BcFV5 (Figure 5C). Two *S. sclerotiorum* strains (A-EV-374-1 and -5) also acquired BcVV1. All six *B. cinerea* derivatives acquired BcVV1, BcFV5, and BcHV5 (Figure 5C). Among the *B. cinerea* derivatives, strains B-EV-374-1, -2, -3, -4, and -6 acquired BcHV5, with three of these (B-EV-374-1, -2, and -4) also acquiring BcFV5. Additionally, five strains acquired BcVV1. These results demonstrate that, on *N. benthamiana* leaves, BcHV5 and BcFV5 can be transmitted via cell-free EVs to both *S. sclerotiorum* and *B. cinerea*.

### Mycoviruses from Strain IBc-374 Confer Debilitation on *S. sclerotiorum*

To assess the impact of mycovirus acquisition on *S. sclerotiorum* phenotype, strain IBc-374 was co-inoculated with strain Ep-1PNA367^R^ on *N. benthamiana* leaves. A total of two independent experiments were conducted. In the first experiment, four derivative strains were isolated; in the second experiment, eighteen derivative strains were isolated (Figure 2M). Phenotypic observations were performed on the four *S. sclerotiorum* derivative strains (A-IBc-374-1, -2, -3, and -4) obtained from the first experiment. The four derivative strains harbored 10 to 11 mycoviruses derived from strain IBc-374. These four subcultures shared nine common mycoviruses: BcHV1, BcMV1, BcMV9, BcMV5, BcUV3, BcOLV1, BcHV5, BcAV1, and BcBNV5. Strain A-IBc-374-1 carried an additional mycovirus, BcHV3. Strain A-IBc-374-2 harbored two additional mycoviruses, BcMV2 and BcHV3. Strain A-IBc-374-3 carried two additional mycoviruses, BcMV2 and BcBEV5. Strain A-IBc-374-4 carried an additional mycovirus, BcMV2 (Figure 6A).

**Figure 6.**
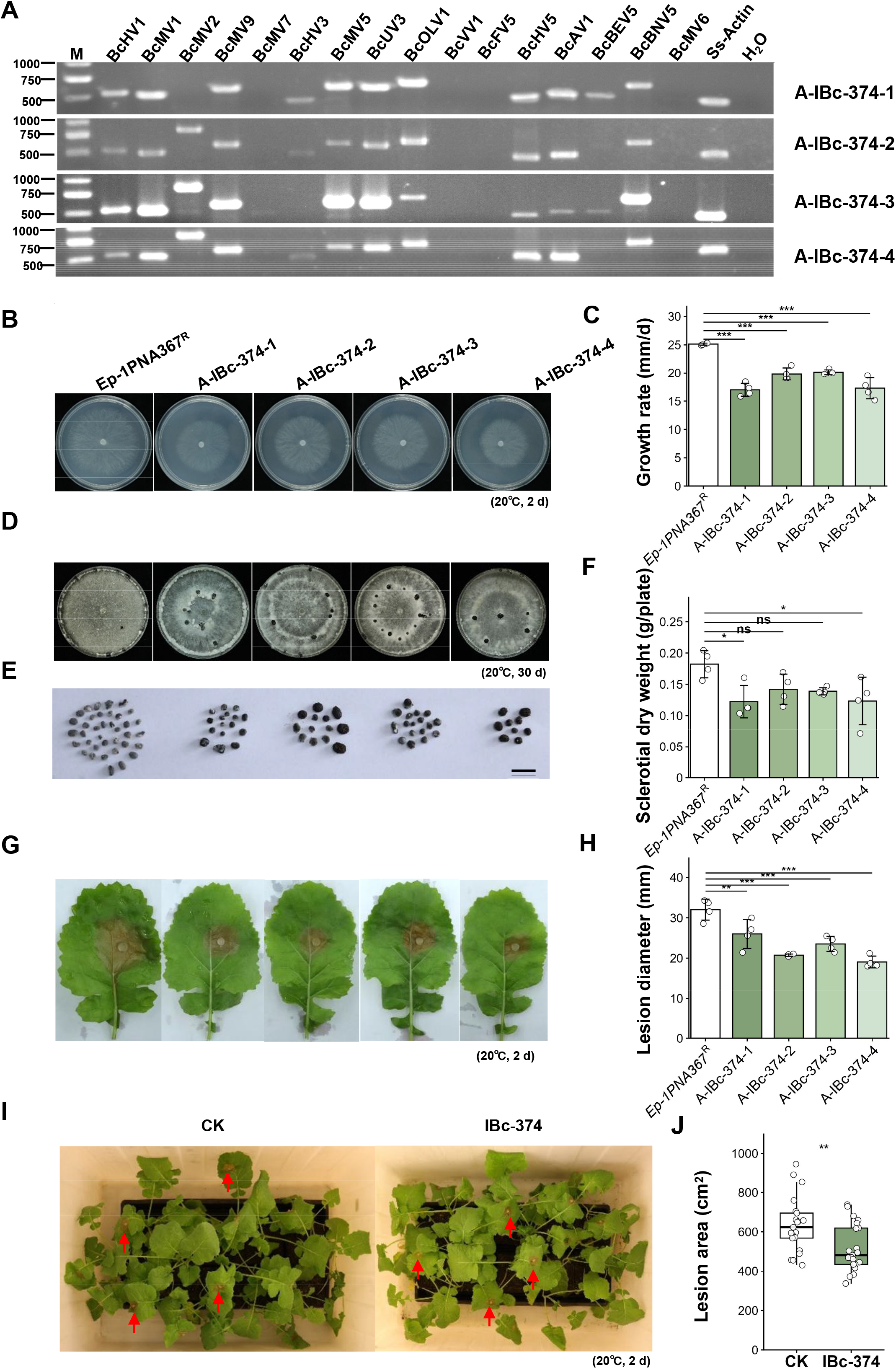
Acquisition of mycoviruses via co-inoculation alters the biological characteristics of recipient *S. sclerotiorum*, and IBc-374 suppresses Sclerotinia stem rot caused by *S. sclerotiorum* on rapeseed. Ep-1PNA367^R^ subcultures were recovered from *N. benthamiana* leaves co-inoculated with IBc-374 and Ep-1PNA367^R^. **(A)** Mycovirus profiles of four representative subcultures examined by RT-PCR. *S. sclerotiorum Actin* was amplified as a positive control for PCR amplification; *B. cinerea tubulin* was amplified as a negative control for PCR amplification. RNA extracted from nuclease-free water were used as negative controls for the RT-PCR template. **(B)** Colony morphology of the four subcultures. Images were taken at 48 hours post inoculation (hpi) on PDA at 20℃. **(C)** Average mycelium growth rate of the four subcultures. Colony diameters were measured at 12 and 24 hpi. Data are from four independent replicates. Error bars indicate standard deviation (SD). Statistical significance was determined by one-way ANOVA with Tukey’s post-hoc test (\*\*\**P* < 0.001). **(D, E)** Sclerotia formation of four subcultures on PDA plates (20℃, 30 days). Scale bar = 1 cm. **(F)** Average sclerotia dry weight of four subcultures. Data are from four independent replicates. Error bars indicate standard deviation (SD). Statistical significance was determined by one-way ANOVA with Tukey’s post-hoc test (\**P* < 0.05, ns *P* ≥ 0.05). **(G, H)** Pathogenicity of the four subcultures on detached rapeseed leaves. **(G)** Representative lesions at 48 hpi (20℃). **(H)** Lesion diameters of the four subcultures on detached rapeseed leaves (20℃, 48 hpi). The experiment was repeated twice. Data are from four independent replicates. Error bars indicate standard deviation (SD). Statistical significance was determined by one-way ANOVA with Tukey’s post-hoc test (\**P* < 0.05, \*\**P* < 0.01, \*\*\**P* < 0.001). **(I, J)** IBc-374 suppresses Sclerotinia stem rot on rapeseed. Rapeseed leaves were pretreated with IBc-374 mycelial suspension, followed by inoculation with Ep-1PNA367^R^ hyphal plugs. H_2_O pretreatment served as a negative control. Plants were maintained at 20℃ under high humidity (100%). **(I)** Disease progression of Sclerotinia rot at 48 hpi. **(J)** Lesion areas measured at 48 hpi. Each sample had twenty-one replicates. The error bar represent standard deviation (SD). Statistical significance was determined by Student’s t test (\*\**P* < 0.01). All experiments were independently repeated twice.

Compared to the parental strain Ep-1PNA367^R^, all four mycovirus-carrying subcultures exhibited significantly reduced radial growth rates on PDA (Figure 6B, C). The average growth rates were 17.0 ± 1.13 mm/d, 19.8 ± 1.08 mm/d, 20.2 ± 0.42 mm/d, and 17.3 ± 1.85 mm/d, respectively, compared to 25.1 ± 0.18 mm/d for Ep-1PNA367^R^ (ANOVA, *P* < 0.001, *n* = 4).

Sclerotia are dormant structures of *S. sclerotiorum*. Under suitable conditions, they can germinate to directly produce infecting hyphae, or germinate to form apothecia that release ascospores. The number of sclerotia in the field is one of the key factors for the epidemic of *Sclerotinia* disease in the next season. We found that the mycovirus-infected *S. sclerotiorum* strains exhibited altered sclerotial production. While all strains produced sclerotia (Figure 6D), the number of sclerotia produced by the four subcultures was significantly lower than that of Ep-1PNA367^R^ (Figure 6E). The dry weights of sclerotia formed by the four subcultures were reduced compared to Ep-1PNA367^R^ (0.18 ± 0.02 g/plate). Specifically, subcultures A-IBc-374-1 (0.12 ± 0.03 g/plate) and A-IBc-374-4 (0.12 ± 0.04 g/plate) showed a statistically significant reduction (ANOVA, *P* < 0.05, *n* = 4), whereas A-IBc-374-2 (0.14 ± 0.02 g/plate) and A-IBc-374-3 (0.14 ± 0.01 g/plate) exhibited a reduction that did not reach statistical significance (Figure 6F).

The virulence of all four subcultures was significantly decreased when inoculated on rapeseed leaves. The average lesion diameters were 26.0 ± 3.58 mm, 20.7 ± 0.29 mm, 23.5 ± 1.85 mm, and 19.0 ± 1.46 mm for strains A-IBc-374-1, A-IBc-374-2, A-IBc-374-3, and A-IBc-374-4, respectively, while the lesion diameter caused by strain Ep-1PNA367^R^ was 32.0 ± 2.59 mm (Figure 6G, H; ANOVA, *P* < 0.05, *n* = 4).

These results demonstrate that the acquisition of multiple mycoviruses from *B. cinerea* strain IBc-374 significantly debilitates *S. sclerotiorum*, reducing hyphal growth, sclerotial formation, and virulence.

### Strain IBc-374 Inhibits *S. sclerotiorum* Infection on rapeseed

To evaluate the biocontrol potential of strain IBc-374 against *S. sclerotiorum* infection, we performed a pre-treatment assay on rapeseed leaves. Pre-treatment with strain IBc-374 hyphal suspension 24 hours prior to challenge inoculation with Ep-1PNA367^R^ significantly suppressed disease progression, reducing lesion area by 18.2% compared to mock-treated controls (519.2 ± 119.86 mm² vs 634.9 ± 140.10 mm² at 48 hpi) (Figure 6I, J; t-test, *P* < 0.01, *n* = 21). Similar to co-inoculation of *S. sclerotiorum* with IBc-374 on *N. benthamiana* (Figure 6A), on rapeseed leaves sprayed with hyphal fragments of IBc-374, *S. sclerotiorum* can also obtain the mycovirus carried by IBc-374, and after virus infection, its growth, pathogenicity, and ability to produce sclerotia are all significantly suppressed (Figure S2).

These results further confirm that the viral transimission can occur both on model plant and field crop, and indicate that intergeneric mycovirus transmission from *B. cinerea* to *S. sclerotiorum* contributes to the biocontrol activity of strain IBc-374, reducing the severity of Sclerotinia rot on rapeseed. The acquisition of multiple mycoviruses leads to debilitation of *S. sclerotiorum*, likely mainly contributing to the observed disease suppression.

### Strain IBc-374 Induces Broad-Spectrum Resistance Against pathogens

To further investigate whether strain IBc-374 can inhibit infection by the highly virulent *B. cinerea*, the mycelial suspension of strain IBc-374 was sprayed onto *N. benthamiana* leaves, followed by inoculation with strain B05.10^G^. The results showed that the lesion area on IBc-374-treated plants (80.4 ± 38.95 mm^2^) was significantly smaller than that on water-treated control plants (450.8 ± 95.81 mm^2^) (Figure 7A, B). From the lesion margins of *N. benthamiana* pretreated with the mycelial suspension of strain IBc-374, four derivative strains of *B. cinerea* B05.10^G^ were recovered. Unlike on PDA no virus can transmit from IBC-374 to *B. cinerea* B05.10^G^, RT-PCR analysis revealed that these derivatives acquired up to 14 mycoviruses, including BcHV1, BcMV1, BcMV9, BcMV7, BcHV3, BcMV5, BcUV3, BcOLV1, BcVV1, BcFV5, BcHV5, BcAV1, BcBNV5, and BcMV6 (Figure 2D). Phenotypic characterization further demonstrated that these virus-carrying *B. cinerea* strains exhibited varying degrees of debilitation, including reduced mycelial growth, attenuated virulence, and decreased sporulation capacity (Figure 2E-K). Collectively, these findings indicate that strain IBc-374 is capable of transmitting mycoviruses not only to *S. sclerotiorum* but also to *B. cinerea* on plants, thereby inducing hypovirulence in both fungal pathogens and exhibiting potential for simultaneous biocontrol of two distinct fungal diseases.

**Figure 7.**
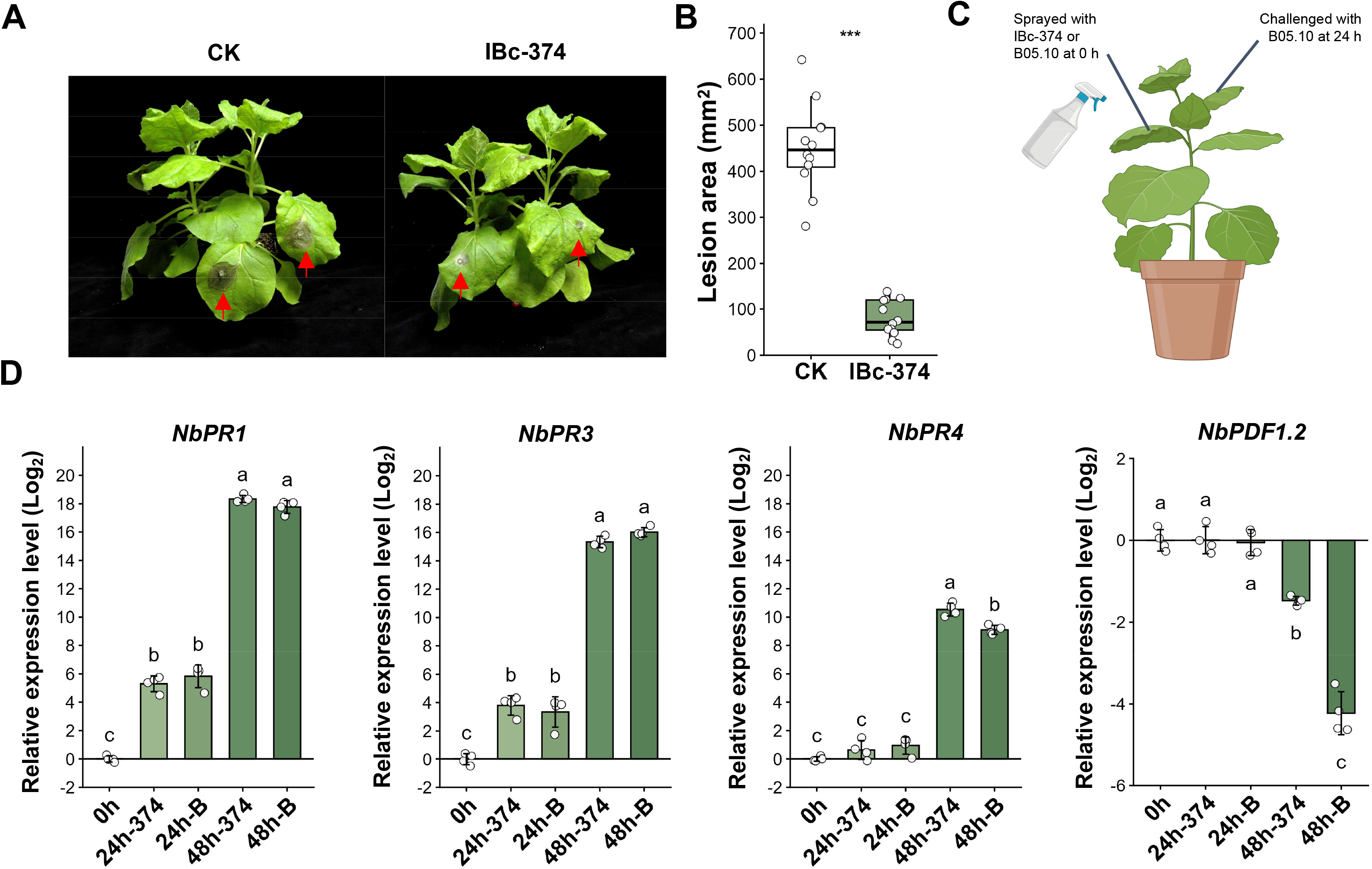
Strain IBc-374 induces plant broad-spectrum resistance against pathogens. **(A)** Disease progression of gray mold on *N. benthamiana*. The hyphal suspension of strain IBc-374 was sprayed on *N. benthamiana*, with water as a control. At 3 days post treatment, B05.10^G^ hyphal plugs were inoculated on *N. benthamiana* leaves at 20℃. **(B)** Lesion area of gray mold on *N. benthamiana*. Error bars indicate standard deviation (SD). Data were analyzed using Student’s *t* test, \*\**P* < 0.01. The experiment was independently repeated twice. **(C)** Schematic diagram of the experimental design for assessing systemic resistance induced by IBc-374. The fifth leaf of the second whorl of 5-week-old tobacco plants was sprayed with hyphal suspension (OD_600_ = 1.2) of strain IBc-374 or B05.10. After 24 h, the upper systemic leaves were challenged by spraying with B05.10 hyphal suspension (OD_600_ = 1.2). Leaf discs were collected from challenged systemic leaves at 0, 24, and 48 h post-challenge. **(D)** Expression levels of tobacco immune-related genes in systemic leaves. Total RNA extracted from leaf discs was analyzed by RT-qPCR. Error bars indicate standard deviation (SD). The experiment was independently repeated twice. Data represent four independent biological replicates. Statistical analysis was performed using ANOVA, followed by Tukey’s post-hoc test, with statistical significance set at *P* < 0.01.

To determine whether strain IBc-374 activates plant immune-related signaling pathways, the fifth leaf of the second whorl of *N. benthamiana* leaves was sprayed with the mycelial suspension of strain IBc-374, using the mycelial suspension of the virus-free, highly virulent strain B05.10 as a control. At 24 hours post-treatment, the upper systemic leaves were challenged by spraying with the mycelial suspension of strain B05.10 (Figure 7C). The results showed that 24 hours after pretreatment with strain IBc-374 on the fifth leaf, the expression of *NbPR1*, a marker gene of the salicylic acid (SA) pathway, as well as the pathogenesis-related protein genes *NbPR3* and *NbPR4*, was significantly activated in the upper systemic leaves. Compared to the 0-hour time point, their expression levels were upregulated by 39.3-fold, 14.0-fold, and 1.5-fold, respectively (Figure 7D). Similarly, pretreatment with the mycelial suspension of strain B05.10 on the fifth leaf of the second whorl for 24 hours also activated the expression of *NbPR1*, *NbPR3*, and *NbPR4* in systemic leaves, reaching transcriptional levels comparable to those induced by strain IBc-374, with no statistically significant difference between the two treatments. At this time point, the expression of *NbPDF1.2*, a marker gene of the jasmonic acid (JA) pathway, did not change significantly.

After pretreatment on the fifth leaf of the second whorl for 24 hours, the upper systemic leaves were challenged with the mycelial suspension of strain B05.10, and the expression of defense-related genes in systemic leaves was examined at 48 hours. Both the IBc-374-pretreated group and the B05.10-pretreated control group exhibited strong upregulation of *NbPR1*, *NbPR3*, and *NbPR4* at the 48-hour time point. Notably, the expression level of *NbPR4* in the IBc-374-treated group was significantly higher than that in the B05.10-treated group. In contrast, the expression of the JA pathway marker gene *NbPDF1.2* was significantly suppressed. Compared to the 0-hour time point, *NbPDF1.2* expression in the upper systemic leaves was downregulated by 2.7-fold in plants pretreated with strain IBc-374, whereas it was downregulated by 20.0-fold in plants pretreated with strain B05.10, indicating that IBc-374 pretreatment effectively alleviates the suppression of *NbPDF1.2* expression.

In summary, both strain IBc-374 and strain B05.10 activated the expression of the SA pathway marker gene *NbPR1* and the downstream defense-related genes *NbPR3* and *NbPR4* in *N. benthamiana*. However, unlike the highly virulent strain B05.10, strain IBc-374 significantly alleviated the pathogen-mediated suppression of *NbPDF1.2*, a JA pathway marker gene, while activating the SA pathway.

## DISCUSSION

In this study, we demonstrate that EVs secreted by the hypovirulent *B. cinerea* strain IBc-374 mediate cross-genus transmission of multiple mycoviruses to *S. sclerotiorum*. Using a combination of cell-free EV incubation, co-culture, and in planta assays, we provide multiple lines of evidence that EVs act as non-cellular vectors for mycovirus spread. To our knowledge, this is the first report that fungal EVs can directly deliver infectious mycoviruses across genus barriers, significantly expanding our understanding of mycovirus ecology and horizontal transfer.

### EV-mediated transmission bypasses hyphal anamostosis

The efficient transmission of mycoviruses from IBc-374 to the intergeneric *S. sclerotiorum* strain Ep-1PNA367^R^ occurred despite clear no hyphal anamostosis between the two fungi at their hyphal interaction zones, as revealed by Evans blue staining (Figure 2B). Notably, *B. cinerea* and *S. sclerotiorum* belong to phylogenetically distinct genera and are incapable of hyphal anastomosis, yet up to 13 mycoviruses were successfully transmitted during dual culture or plant co-inoculation (Figure 2). When cell-free EVs were incubated with protoplast-derived young hypae of Ep-1PNA367^R^, successful virus acquisition occurred (Figure 4D). This demonstrates that *S. sclerotiorum* is intrinsically capable of taking up EVs and supporting mycovirus replication independently of hyphal anastomosis. At least two mechanisms for viral transimisson between two vegetative incompatible strains of *B. cinerea* in planta, the one is via EVs because *B. cinerea* can acquire mycovires from cell-free EVs, while another is by weakening vegetative incompatiblity reaction through proline scereted by *B. cinerea*-infected plant^12^. Thus, EV-mediated delivery effectively uncouples virus transmission from hyphal anastomosis and can overcome vegetative incompatibility among strains of a species, a major bottleneck in mycovirus biocontrol.

### Selective packaging and transmission efficiency

Among the 16 mycoviruses in IBc *-* 374, only a subset (4 *-* 7) was consistently detected in purified EV preparations (Figure 3E), and even fewer successfully established infection in recipient fungi after EV feeding (Figure 4F). Notably, BcMV1, the most abundant intracellular virus, was rarely detected in EVs (5.3% frequency), whereas BcHV1, one of the least abundant, was detected more frequently (15.8%). Interestingly, further analysis revealed that certain viruses-namely BcHV5, BcFV5, BcBNV5, and BcHV1-exhibited a consistent positive correspondence between their intracellular abundance and EV detection frequency (Figure 3I). This observation supports a passive or stochastic encapsulation model for this subset of viruses, where higher cytoplasmic abundance increases the likelihood of random incorporation into EVs during vesicle formation.

In contrast, BcMV1, BcOLV1, and BcUV3 deviated from this trend, with BcMV1 showing the most extreme discordance (highest abundance, lowest EV frequency). These discrepancies suggest that additional virus-specific factors may interfere with the passive packaging process. For instance, BcMV1, a mitovirus, is localized within mitochondria, which may physically sequester its RNA from the EV biogenesis machinery. Similarly, BcOLV1, an ourmia-like virus, also exhibits mitochondrial enrichment of viral RNAs, as previously reported^43^, potentially contributing to reduced EV packaging efficiency. These findings highlight that subcellular localization and distinct replication strategies are important determinants of viral encapsidation into EVs.

Despite the differential packaging patterns observed, the discrepancy between detection in EVs and successful transmission (e.g., BcHV1 was detected in EVs but not transmitted in the EV-feeding experiment) indicates that EV packaging alone is insufficient. BcHV1 and BcBNV5 failed to be transmitted to recipient strains, likely attributable to their low intracellular abundance (Figure 1F), which probably results in low copy numbers in EVs. Additional factors such as viral integrity, copy number, co-packaging of necessary replication machinery, or recipient cell compatibility likely determine productive infection. Furthermore, given that some mycoviruses have defined host ranges, the possibility that virus-recipient incompatibility also contributes to the lack of transmission cannot be ruled out. Future studies using quantitative RT-PCR or RNA-seq on density-gradient-purified EVs are needed to precisely measure packaging efficiency and to identify potential RNA motifs or host proteins that govern selective sorting.

### The curious case of BcVV1

The victorivirus BcVV1 was not protected by EV membranes in nuclease protection assays (Figure 3F), suggesting it is not encapsulated inside EVs. Nevertheless, BcVV1 was transmitted to recipient fungi in both EV-feeding and in planta experiments (Figure 5C), and was detected in recipient strains of both fungal species, possibly due to its strong infectivity^30^. This implies that our *“*EV-enriched fraction*”* likely contains co-purified viral particles or other non-EV nanoparticles. The successful transmission of BcVV1, despite its existence as free viral particles rather than being encapsulated within EVs, suggests that alternative, EV-independent mechanisms may also contribute to virus dissemination. This does not diminish the main conclusion, EVs clearly transmit several other mycoviruses, but highlights the need for more stringent EV purification (e.g., density gradient ultracentrifugation) in future mechanistic studies. The co-existence of EV-encapsulated and free viral particles in the same preparation may even reflect a natural scenario where both modes operate synergistically.

### Recipient fungi acquire more mycoviruses than those detected in purified EVs

In this study, the number of mycoviruses transmitted to recipient fungi consistently exceeded those detected in purified EVs. For example, while only 7 mycoviruses were detected in EV preparations, up to 13 were transmitted to *S. sclerotiorum*. Several possibilities may explain this discrepancy. First, some viral RNAs may be present in EVs at levels below RT-PCR detection limits yet remain infectious. Second, certain mycoviruses may be transmitted through EV-independent routes, as exemplified by BcVV1, which was not encapsulated in EVs but was still transmitted. Third, upon entering recipient cells, low-level viral RNAs may replicate to readily detectable levels. Additionally, our EV purification may not capture all EV subpopulations or may underestimate the full cargo diversity. Thus, EVs likely serve as a major but not exclusive vehicle for mycovirus transmission, and the coexistence of multiple pathways may ensure robust viral spread across fungal species.

### Implications for fungal ecology

The discovery of EV-mediated cross-genus transmission provides a plausible explanation for the long-standing observation that identical or highly similar mycoviruses are found in phylogenetically divergent fungi ^34,35^. EVs may serve as “mobile virus reservoirs” that protect viral RNA/DNA from environmental degradation ^25^ and facilitate spread across species boundaries without requiring direct cell-to-cell contact. This mechanism could be particularly relevant in soil or phyllosphere communities, where hyphae of different fungal taxa are in close proximity but typically do not undergo anastomosis^44^. Future environmental metatranscriptomic studies, coupled with EV isolation from complex microbial communities, could test the broader ecological significance of this pathway. Specifically, in the case of *B. cinerea* and *S. sclerotiorum*, EV-mediated mycovirus transmission from the former to the latter strongly impacts the survival of *S. sclerotiorum* by reducing its virulence, stunting its growth, and decreasing sclerotial production, thereby weakening its competitive ability agains*t B. cinerea* which shares the same ecological niche.

### Dual functions for disease control: virus transmission plus plant immunity

The superior biocontrol activity of strain IBc-374 or its EV-rich hyphal suspension, as evidenced by significant disease reduction on both *N. benthamiana* and rapeseed (Figure 6, 7), likely stems from two complementary mechanisms that operate in concert. First, it directly disarms the pathogen by transferring hypovirulenc-associated mycoviruses via EVs (and possibly free viral particles), leading to the debilitated phenotypes observed in re*-*isolated *S. sclerotiorum* and *B. cinerea* strains (Figure 2, Figure 6, and Figure S2). Second, it fortifies the host by priming plant immunity: treatment with the hyphal suspension up-regulated defence-related genes including NbPR1, NbPR3, and NbPR4 (Figure 7D), indicative of salicylic acid (SA) pathway activation ^45^. Notably, unlike the virulent strain B05.10, IBc-374 treatment alleviated the suppression of the jasmonic acid (JA) pathway marker NbPDF1.2, suggesting a more balanced immune activation that may contribute to its broad-spectrum efficacy. While these gene expression data point to induced resistance, we acknowledge that functional proof, e.g., using chemical inhibitors or plant signalling mutants-is required to firmly establish the necessity of SA or JA signalling in the observed protection. Nevertheless, this *‘* disarm and fortify *’* dual action makes IBc *-* 374 a particularly attractive biocontrol agent against two devastating diseases that often co-occur in the field ^46,47^, and highlights the potential of EV*-*mediated virus delivery as a strategy for sustainable disease management.

### Towards EV-based delivery platforms for mycoviruses

The ability of cell-ree EVs to deliver infectious mycoviruses to protoplast-derived germlings (Figure 4) and to fungi in planta (Figure 5) raises the possibility of developing EV-mimetic or synthetic delivery systems. For instance, hypovirulence-associated viral RNAs could be encapsulated in liposomes or engineered EVs for direct application as “mycovirus-derived RNA pesticides”. The recent regulatory approval of the RNA-based pesticide Ledprona (Calantha^Ⓡ^)^48^ underscores the feasibility of such approaches. Key challenges ahead include optimizing EV production yield, achieving high-efficiency loading of selected viral genomes, and ensuring stability under field conditions. Nevertheless, our study provides a proof of concept that EV-mediated transmission can be harnessed for biological control, opening a new frontier in mycovirus application.

### Limitations and future directions

Several questions remain unresolved. First, although EV secretion has been documented in both unicellular and multicellular fungi^15,38,39,49^, the biogenesis pathway of EVs loaded with mycoviruses remains unknown. Does the ESCRT machinery participate in this process^50–54^ Do different viruses exploit distinct EV subpopulations? Notably, in this study,the mycoviruses transmitted via EVs are capsidless; Second, what determines the host range of EV-mediated transmission? The same EV preparation infected *S. sclerotiorum* and *B. cinerea* germlings with different efficiencies (e.g., BcHV5 was more efficient in *B. cinerea*, BcOLV1 was comparable in both), suggesting species-specific factors influence viral entry or replication. Third, the relative contribution of EV-mediated versus direct hyphal transmission in natural settings awaits quantification. Answering these questions will require a combination of genetics (knockout of predicted EV biogenesis genes), proteomics (cargo identification), and advanced imaging.

In conclusion, our study identifies fungal extracellular vesicles as a previously unrecognized vehicle for cross-genus mycovirus transmission, overcomes the long-standing barrier of vegetative incompatibility, and demonstrates a dual-action biocontrol strategy against two major fungal pathogens. These findings not only advance fundamental knowledge of viral ecology but also lay the groundwork for EV-based mycovirus delivery platforms in sustainable agriculture.

## METHODS

### Fungal strains, culture conditions and biological characterization test

*B. cinerea* strain IBc-374 was isolated from strawberry in 2019^55^. The highly virulent, mycovirus-free *B. cinerea* strain B05.10 and its neomycin-resistant derivative B05.10^G^ ^56,57^ were used in this study. For *S. sclerotiorum*, the highly virulent strain Ep-1PNA367^58^ and its hygromycin B-resistant derivative Ep-1PNA367^R^ ^12^ were employed. All strains were cultured on potato dextrose agar (PDA) at 20℃ and stored at 4℃.

The hyphal growth rate and pathogenicity were determined as previously described^56^. At 30 days post-inoculation (dpi), the sclerotia produced by *S. sclerotiorum* strains on PDA were harvested, air-dried, and subsequently quantified by gravimetric analysis. Digital images were captured using a Canon PC1817 digital camera (Canon Inc., Tokyo, Japan).

Conidiation of *B. cinerea* was induced on MEM (Malt Extract Medium) at 20℃. After 15 days of incubation, the sporulation of each strain was examined. The conidia were harvested by washing each plate with 5 mL of sterile distilled water, and the spore concentration was determined using a hemocytometer. The total spore yield per plate was then calculated based on the volume.

To collect EVs from *B. cinerea* strains, the fungal strains were inoculated in rapeseed leaf medium and incubated statically at 20℃ for 4 days. The culture filtrate was then collected for vesicle extraction. The rapeseed leaf medium was prepared as follows: 10 g of fresh rapeseed leaves were mixed with distilled water and homogenized using a juicer. Subsequently, 2 g K_2_HPO_4_, 1.45 g KH_2_PO_4_, 0.6 g MgSO_4_**·**7H_2_O, 0.3 g NaCl, 0.001 g FeSO_4_, 0.5 g (NH_4_)_2_SO_4_, and 0.01 g CaCl_2_**·**2H_2_O were added. The volume was brought to 1000 mL with distilled water, the pH was adjusted to 7.0, and the medium was sterilized by autoclaving at 121℃ for 20 min.

### RNA extraction and mycovirus detection

Total RNA was extracted using TRIzol reagent (Diyue Biotechnology, Wuhan, China). Gel electrophoresis and NanoDrop 2000 (Thermo Fisher Scientific Inc.) were used to detect the quality and concentration of the RNA. The complementary DNA (cDNA) was reverse transcribed from extracted total RNA using a cDNA synthesis kit (Transgen Biotech, Beijing, China). Mycoviruses were detected using mycovirus-specific primers by reverse transcription polymerase chain reaction (RT-PCR).

To detect the relative viral content in strain IBc-374, real-time PCR was performed using mycovirus-specific primers (amplifying fragments ranging from 100 bp to 165 bp). The real-time PCR amplification was carried out using the PerfectStart^®^ Green qPCR SuperMix kit. A three-step qPCR procedure was performed as follows: initial denaturation at 94℃ for 30 seconds, followed by 40 cycles of denaturation at 94℃ for 5 seconds, annealing at 57℃ for 15 seconds, and extension at 72℃ for 10 seconds. Fluorescence signals were measured at the end of each extension step. The *Bcactin* gene was used as an internal reference. Relative viral RNA levels were calculated using the 2^-ΔΔCt^ method, and log_2_-transformed values were used for statistical analysis to normalize data distribution. Each virus was assayed in five independent biological replicates. The primers used in this study are listed in Table S2.

EVs were isolated from the culture filtrate of IBc-374. Total RNA was extracted from the purified EVs using TRIzol reagent. The presence of each mycovirus in the EVs was then detected by RT-PCR using virus-specific primers. The frequency of detection for each mycovirus was calculated based on 19 independent experimental replicates, and the frequency of each mycovirus was calculated.

### Detection of mycoviruses by Northern blot

To detect BcHV5, Northern blot analysis was performed. A DNA probe specific to BcHV5 was amplified by PCR using BcHV5-specific primer pairs. The probe was then labeled with digoxigenin (DIG) using the DIG-High Prime DNA Labeling and Detection Starter Kit I (Roche Diagnostics GmbH, Mannheim, Germany) according to the manufacturer’s instructions. Total RNA extracted from either fungal mycelia or purified EVs was separated by denaturing agarose gel electrophoresis and subsequently transferred onto a positively charged nylon membrane (Amersham Hybond-N+, GE Healthcare) by capillary transfer. The RNA was crosslinked to the membrane by UV irradiation. The membrane was then prehybridized and hybridized with the DIG-labeled probe at 28℃. Following hybridization, the membrane was washed under stringent conditions. Hybridized signals were detected using anti-digoxigenin-AP conjugate and NBT/BCIP substrate, producing purple-blue precipitates. The membrane was air-dried and photographed using a ChemiDoc Touch Imaging System (Bio-Rad). For controls, total RNA extracted from strain IBc-374 was used as a positive control, and total RNA from strain B05.10 was used as a negative control.

### Mycovirus horizontal transmission assay

To clarify the characteristics of horizontal transmission of the 16 mycoviruses on PDA, the donor strain IBc-374 was dual-cultured with the receptor *S. sclerotiorum* strain Ep-1PNA367^R^, as previously described^59^. After the mycelia of the donor and receptor strains came into contact sufficiently for 7 days, hyphal plugs were taken from the receptor strain in the area away from the donor strain IBc-374 and inoculated onto PDA containing the corresponding antibiotic (hygromycin B 50 μg/μL) for cultivation.

The transmission characteristics of these mycoviruses were also investigated in planta. The donor strain IBc-374 was co-inoculated with receptor strain Ep-1PNA367^R^ on leaves of tabaco (*N. benthamiana*). Then, the plants were incubated under alternating light and dark conditions (12 h light and 12 h dark) for 2-4 days. The receptor strains were rescued from lesions and inoculated on PDA amended with both 100 μg/μL cephalosporin and 50 μg/μL hygromycin B.

After three successive subcultures, mycoviruses in the recipient strain Ep-1PNA367^R^ were detected by RT-PCR. The experiment was conducted twice, yielding 26 strains isolated from PDA and 22 strains isolated from plant lesions. These isolates originated from strain Ep-1PNA367^R^.

### Evans blue staining

To assess vegetative compatibility responses between fungal strains, Evans blue staining was employed. Strains IBc-374 and Ep-1PNA367^R^ were inoculated at opposite ends of a PDA covered with cellophane. After 20 hours of contact, they were stained for 30 min at 20℃ using a 0.5% (w/v) Evans blue solution. After washing twice with phosphate buffer saline (PBS, pH = 7.4), the contact area of two colonies was examined to determining their vegetative compatibility reaction.

### EV isolation and nanoparticle tracking analysis (NTA)

The isolation of EVs was performed as described by Cai et al.^60^, with minor modifications. To prevent contamination by intracellular vesicles, EVs were collected from the culture filtrate using a stationary liquid culture method. *B. cinerea* strains were incubated statically in rapeseed leaf medium for 4 days at 20℃. The culture supernatant was subsequently subjected to sequential centrifugation at 4℃ as follows: 300 g for 10 min, 2,000 g for 10 min, and 10,000 g for 30 min. The resulting supernatant was filtered through a 0.45 µm filter membrane and then centrifuged at 100,000 g for 60 min at 4℃. The EVs were washed with 1 × PBS and centrifuged again at 100,000 g for 1 h. The PBS used for washing and resuspending EVs was pre-cleared by ultracentrifugation at 100,000 × g for 60 min at 4 ℃.

The number and size of the EVs were measured using NTA by a ZetaView Particle Metrix (Particle Metrix, PMX-120, Germany). The instrument was calibrated with 100 nm polystyrene beads (Thermo Fisher Scientific, Fremont, CA) prior to each measurement. The washed EV pellet was resuspended in pre-cleaned PBS to form the EV suspension. The final EV suspension was then diluted with pre-cleaned PBS to an appropriate concentration (approximately 1 × 10⁸ particles/mL) for NTA measurement. Three 60-second videos were recorded per replicate at 25℃. The NTA software was used to automatically capture and analyze particle tracks. The batch processing function was applied to calculate the final nanoparticle concentration (particles/mL) and median size (nm).

### Ultrastructural observation of EVs by electron microscopy

To observe the internal structure of hyphae, fresh hyphae were processed according to Cai et al.^60^. Ultrathin sections were collected and stained with uranyl acetate and lead citrate. The sections were observed using a transmission electron microscope (TEM) at an accelerating voltage of 200 kV.

For ultrastructural characterization of EVs, 10 μL of EV suspension were placed onto a copper grid for 5 min. Phosphotungstic acid solution was then added to the copper grid for 5 min. The samples were air-dried and observed under TEM.

### Nuclease protection assay

To determine whether mycoiruses are encapsulated in EVs, the collected EVs, with or without Triton X-100 perforation treatment, were incubated with Micrococcal Nuclease (MNase, Beyotime Biotechnology, Shanghai, China) to assess whether EVs can protect viral nucleic acids from MNase degradation. For Triton X-100 treatment, 200 μL of EV suspension was incubated with 1% Triton X-100 on ice for 30 min. Then, 10 U of MNase was used to treat EV suspension at 37℃ for 15 min. Total RNA from treated EV suspension was extracted for virus detection^61^.

Since the virus BcVV1, which encodes coat protein, was found to be insensitive to perforator treatment in the nuclease (MNase) and perforator (Triton X-100) treatment assays, this suggests that it was most likely collected in the form of viral particles. Therefore, additional experiments involving nuclease, perforator, and proteinase K treatments were designed to further elucidate whether the viral particles of this virus are encapsulated within EVs. After extracting EVs from the culture filtrate, 250 μL aliquots of EVs were subjected to enzymatic treatments under four conditions: (i) untreated as a control; (ii) treated with MNase (10 U); (iii) treated with 1% Triton X-100 on ice for 30 min followed by MNase (10 U); (iv) treated with Proteinase K (100 μg/mL) at 37℃ for 60 min, followed by PMSF to terminate the reaction, and then treated with MNase (10 U). Following the treatments, RNA was extracted from each sample and subjected to RT-PCR using mycovirus-specific primers to detect mycovirus. To determine whether the collected EVs were contaminated with cytoplasmic contents, a highly expressed transcriptional regulator gene, *B. cinerea Sterol Regulator* (*BcSR*, BCIN_02g03500), was used as a negative control, while mycovirus BcHV5 lacking viral particles served as a positive control.

### Co-localization analysis of viral RNA and EVs by fluorescence in situ hybridization (FISH)

Since BcHV5 was detected most frequently in EVs, it was selected as a representative for confirming the co-localization of the viral nucleic acids with EVs by FISH. To prepare the probe, a 566-bp dscDNA fragment of viral genome was amplified using primers BcHV5-probeF/R (Table S2). After purification, the DNA fragment was labeled with Biotin-14-dCTP by the random primer method (BioPrime DNA Labeling System, Invitrogen) and used as the probe. FISH was performed as previously described^62^ with minor modifications. Briefly, after hybridization with the biotin- labeled probe, the EV samples were incubated with Cy5- conjugated streptavidin (Thermo Fisher Scientific, Fremont, CA, USA) to detect the probe. Then, the phospholipid bilayers of the EV samples were stained with FM4-64 dye (Thermo Fisher Scientific, Fremont, CA) and washed with 1 × PBS. Fluorescent signals were observed under a confocal microscope (Leica SP8, Oskar-Barnack, Germany). FM4-64 was excited at 561 nm with an offset of −10, and its emission was collected within a detection window of 570-616 nm. Cy5 was excited at 638 nm with an offset of 0, and its emission was collected within a detection window of 633-738 nm. To minimize spectral crosstalk, a negative offset was applied to the FM4-64 channel to suppress its long-wavelength emission tail. Spectral unmixing was subsequently applied to separate the two fluorophores.

### Mycovirus transmission via EVs

To determine whether EVs carrying mycoviruses could directly transmit to *B. cinerea* and *S. sclerotiorum in vitro*, EVs were collected from the culture filtrate of strain IBc-374 and incubated with protoplast-derived germlings of receptor strains Ep-1PNA367^R^ and B05.10^G^, respectively. Nanoparticles collected from plain medium served as a negative control. The EVs and the negative control nanoparticles were stained with FM4-64, washed three times with PBS by ultracentrifugation (100,000 g for 60 min at 4℃ per wash), and then incubated with an equal volume of protoplast-derived germlings of receptor strains for 4 h at 20℃. Uptake of stained EVs by the protoplast-derived germlings was observed using a confocal microscope. After co-incubating the protoplast-derived germlings with EVs collected from culture filtrate or nanoparticles collected from blank plain medium, the incubated germlings were washed twice with 1 × PBS buffer, then inoculated onto PDA. Following subculturing, they were transferred onto cellophane-overlaid PDA plates, and the mycelia were subsequently harvested to detect the presence of mycoviruses in the resulting colonies.

To determine the roles of EVs in mycoviral transmission in plants, the purififed EVs were injected into tobacco leaves, followed by inoculation with strains B05.10^G^ and Ep-1PNA367^R^, respectively. The extracted EVs were dissolved in 1 mL of PBS, then diluted 30-fold to adjust the concentration to about 2.0 × 10^11^ particles/mL. Then, 150 μL of EVs were injected into leaves of tobacco plants grown for 5 weeks. B05.10^G^ or Ep-1PNA367^R^ was inoculated at the injection site. After 72 hours incubation, two pathogen-derived strains were isolated from the junction between diseased and healthy tissues, and the virus carriage status of the derived strains was examined. Each treatment had three replicates.

### Hyphal suspension of strain IBc-374 protects against two fungal diseases challenge

To investigate the disease control potential of hyphal suspensions rich in EVs, a 4-day culture of strain IBc-374 was homogenized and adjusted to an OD_600_=1.0. The hyphal suspension was evenly sprayed onto the leaf surfaces of plants, which were then incubated in a moist chamber under a 12-h light/dark cycle for 24 h (water treatment served as the control). Subsequently, the leaves were challenged with mycelial plugs of the pathogens under maintained humid conditions. To evaluate the inhibitory effect on gray mold, 12 pretreated tobacco plants were inoculated with B05.10^G^, and the control efficacy was assessed after 72 hours. To determine the suppressive activity against Sclerotinia stem rot, 21 pretreated rapeseed plants were inoculated with strain Ep-1PNA367^R^, and lesion diameters were measured 48 hpi. Strains Ep-1PNA367^R^ and B05.10^G^ were re-isolated from the lesions for mycovirus detection and biological characterization.

### Quantification of immunity-related gene expression in tobacco

To investigate whether the hyphal suspension of strain IBc-374 could enhance plant immunity, the fifth leaf of the second whorl of 5-week-old tobacco leaves was sprayed with hyphal suspensions (OD_600_ = 1.2) of either strain IBc-374 or B05.10. At 24 hours post-treatment, the upper systemic leaves were challenged by spraying with a hyphal suspension of strain B05.10 (OD_600_ = 1.2). Leaf discs were collected from the challenged systemic leaves at 0, 24, and 48 hours post-spraying. Total RNA was extracted from these leaf discs to examine the expression levels of *N. benthamiana* immune-related genes. The primer information is listed in the Table S2.

### Statistical analysis

All statistical analyses were performed using R (version 4.5). All graphs were generated using R (version 4.5) with the ggplot2 package, with error bars representing standard deviation (SD). All experiments were repeated with the replication numbers indicated in the figure legends. A *P*-value < 0.05 was considered statistically significant.

For data with a normal distribution and homogeneity of variance, two-group comparisons were assessed using two-tailed Student’s t-test; for data that did not meet these assumptions, the Mann-Whitney U test was employed. Differences in intracellular abundance between EV-detected (n = 7) and non-detected viruses (n = 10) were assessed using the Mann-Whitney U test. Spearman’s rank correlation coefficient (ρ) was calculated to examine the relationship between intracellular abundance and EV detection frequency for all 16 viruses (assigning 0% frequency to non-detected viruses).

For transmission analysis, Fisher’s exact test compared transmission rates of four viruses between *S. sclerotiorum* (n = 32) and *B. cinerea* (n = 33) recipients. Logistic regression (binomial model) assessed the effects of virus identity and recipient species on transmission success, with Wald tests for predictor significance. A likelihood ratio test compared the main-effects and interaction models.

For multiple group comparisons, one-way ANOVA with Tukey’s post-hoc test was used.

## ACKNOWLEDGMENTS

This research was supported by the National Natural Science Foundation of China (32130087) and Fundamental and Interdisciplinary Disciplines Breakthrough Plan of the Ministry of Education of China (JYB2025XDXM703).

## Figure legends

**Figure S1 Phylogenetic tree of mycoviruses based on RdRp amino acid sequences.** Viral RdRp amino acid sequences were aligned and used to construct the phylogenetic tree using the maximum-likelihood method with MEGA X. Bootstrap support values based on 1,000 replicates are shown at the nodes.

**Figure S2 Horizontal transmission of IBc-374 mycoviruses to *S. sclerotiorum* via sprayed mycelial suspension on rapeseed alters the biological characteristics of the recipient, related to Figure 6I.** Ep-1PNA367^R^ hyphal plugs were inoculated onto rapeseed leaves after pretreatment with IBc-374 hyphal suspension. Plants were maintained at 20℃ under high humidity (100%). H_2_O was used as a negative control. Ep-1PNA367^R^ subcultures were rescued from lesions of inoculated rapeseed leaves. **(A)** Mycoirus profiles of subcultures examined by RT-PCR. *Actin* of *S. sclerotiorum* was used as a positive control for PCR amplification, and H_2_O was used as a negative control for the PCR template. **(B)** Colony morphology of four representative subcultures. Images were taken at 48 hpi on PDA at 20℃. **(C)** Average mycelial growth rate of four subcultures. Colony diameters were measured at 12 and 24 hpi. Growth rate of each strain was calculated. **(D)** Sclerotia formation of four subcultures on PDA plates after 30 days at 20℃. **(E, F)** Pathogenicity of the four subcultures on detached rapeseed leaves. **(E)** Representative lesions at 48 hpi (20℃). **(F)** Lesion area of four subcultures on detached rapeseed leaves (20℃, 48 hpi). Strains designated P24-1, P24-2, P24-3, and P24-4 were *S. sclerotiorum* isolates recovered from plants that had been sprayed with the IBc-374 mycelial suspension. The strain designated P24-CK was isolated from control plants sprayed with water. Each strain had four replicates and the experiment was repeated twice. Error bars indicate standard deviation (SD). The data were analyzed using Dunnett’s test (\*\**P* < 0.01; \*\*\**P* < 0.001; ns not significant).

**Table S1.**
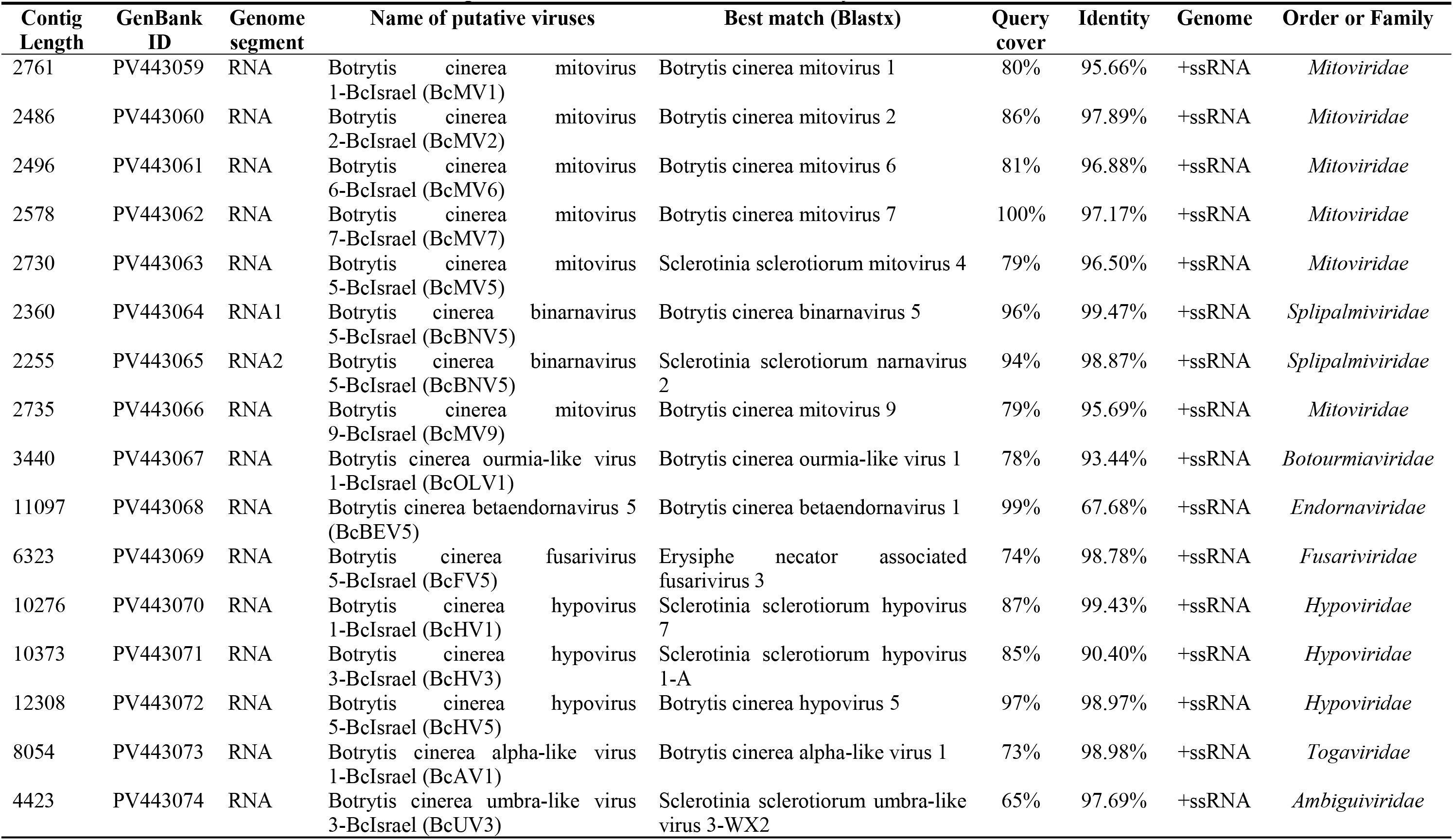
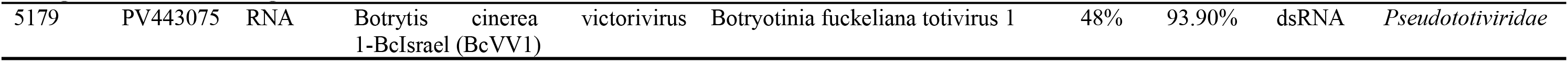
Sequence information of viruses in *Botrytis cinerea* strain IBc-374.

**Table S2.**
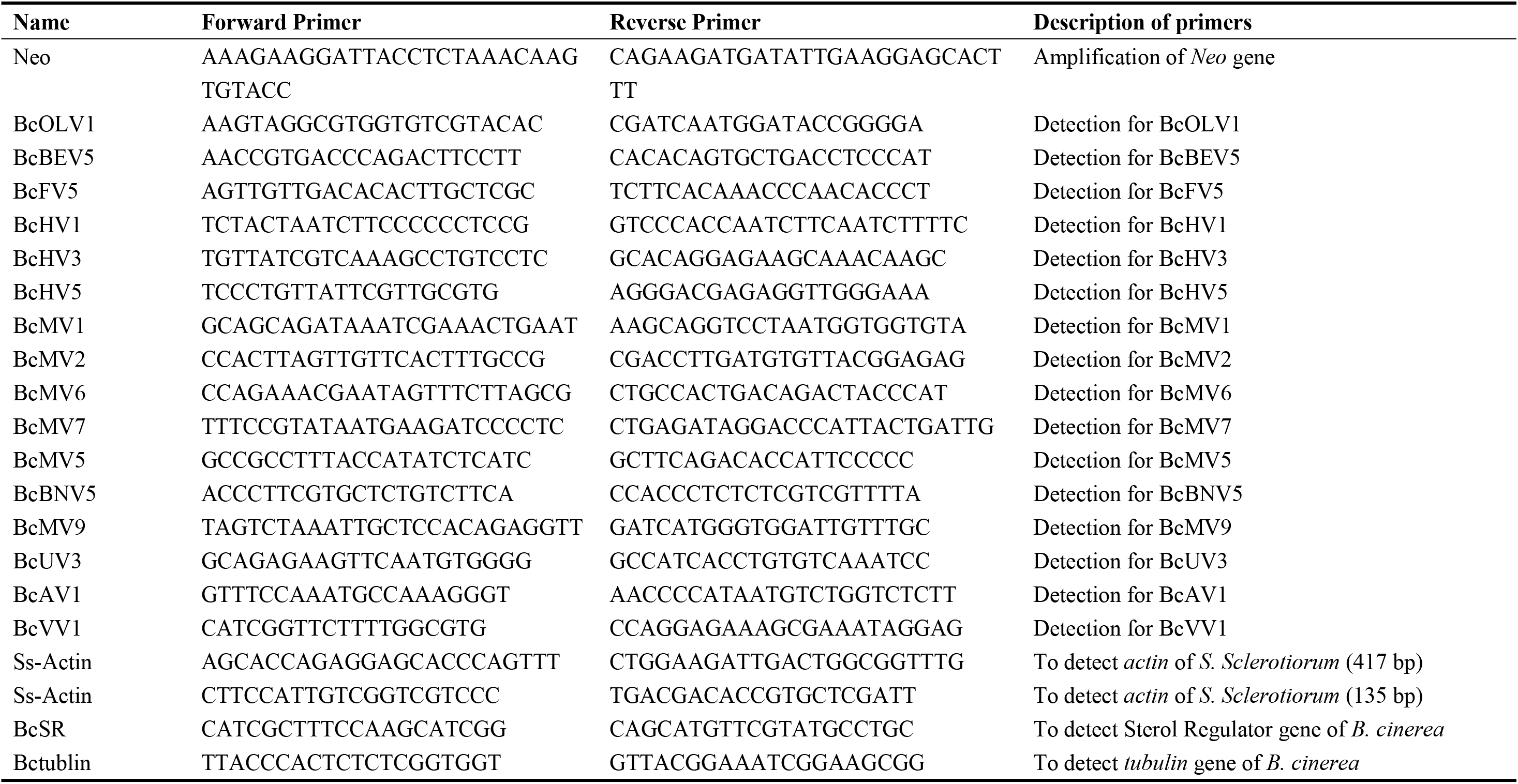
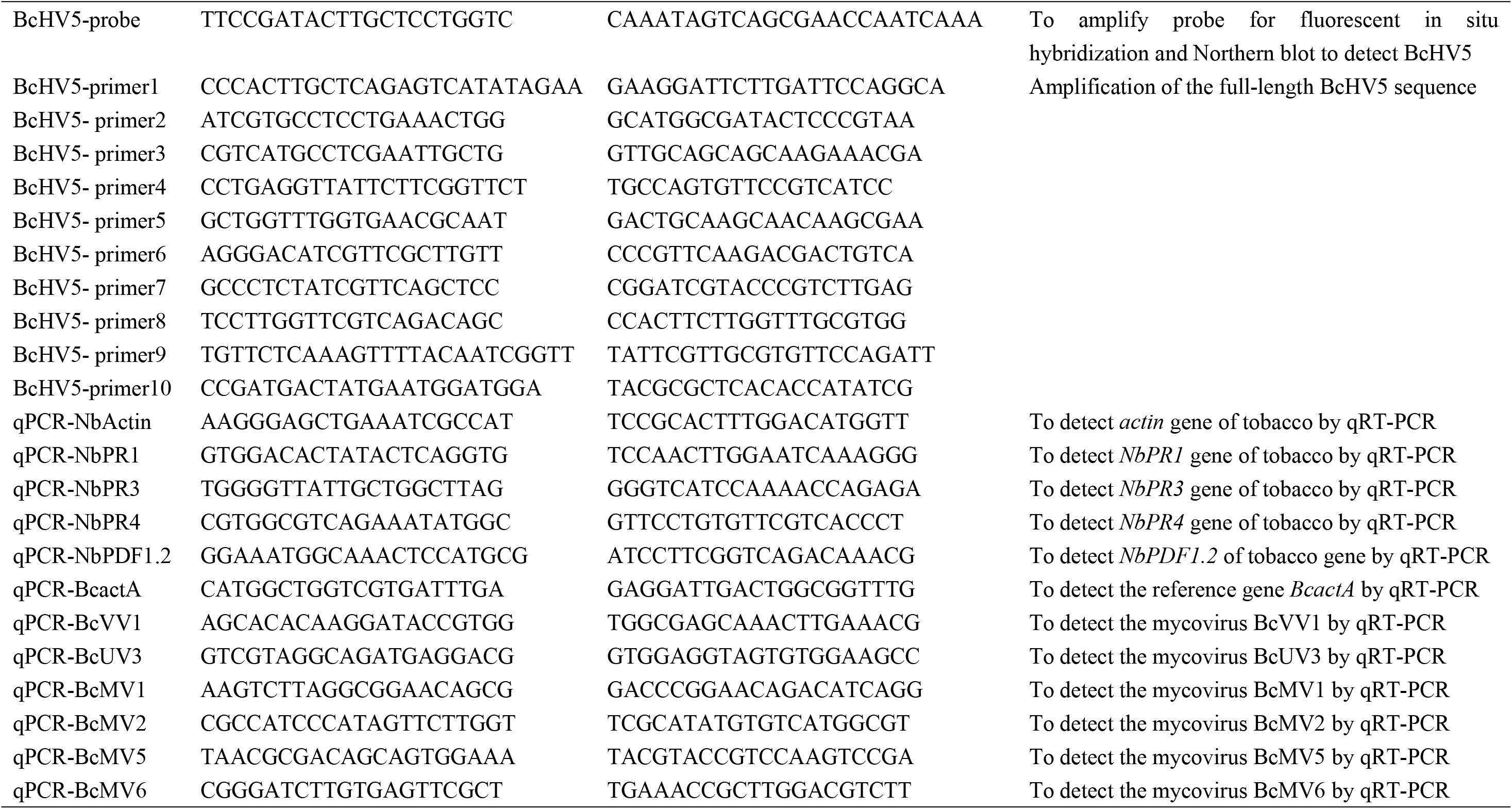
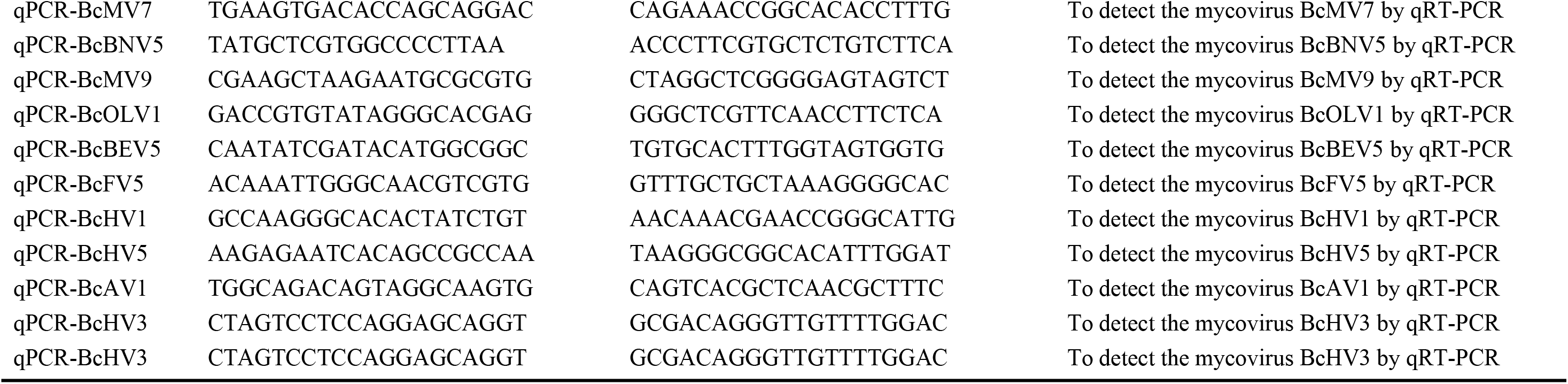
Primers used in this study.

