## Supplementary figures and images for "Extracellular vesicles drive cross-genus mycovirus transmission and suppress two fungal diseases"

### Supplemental Figure S1

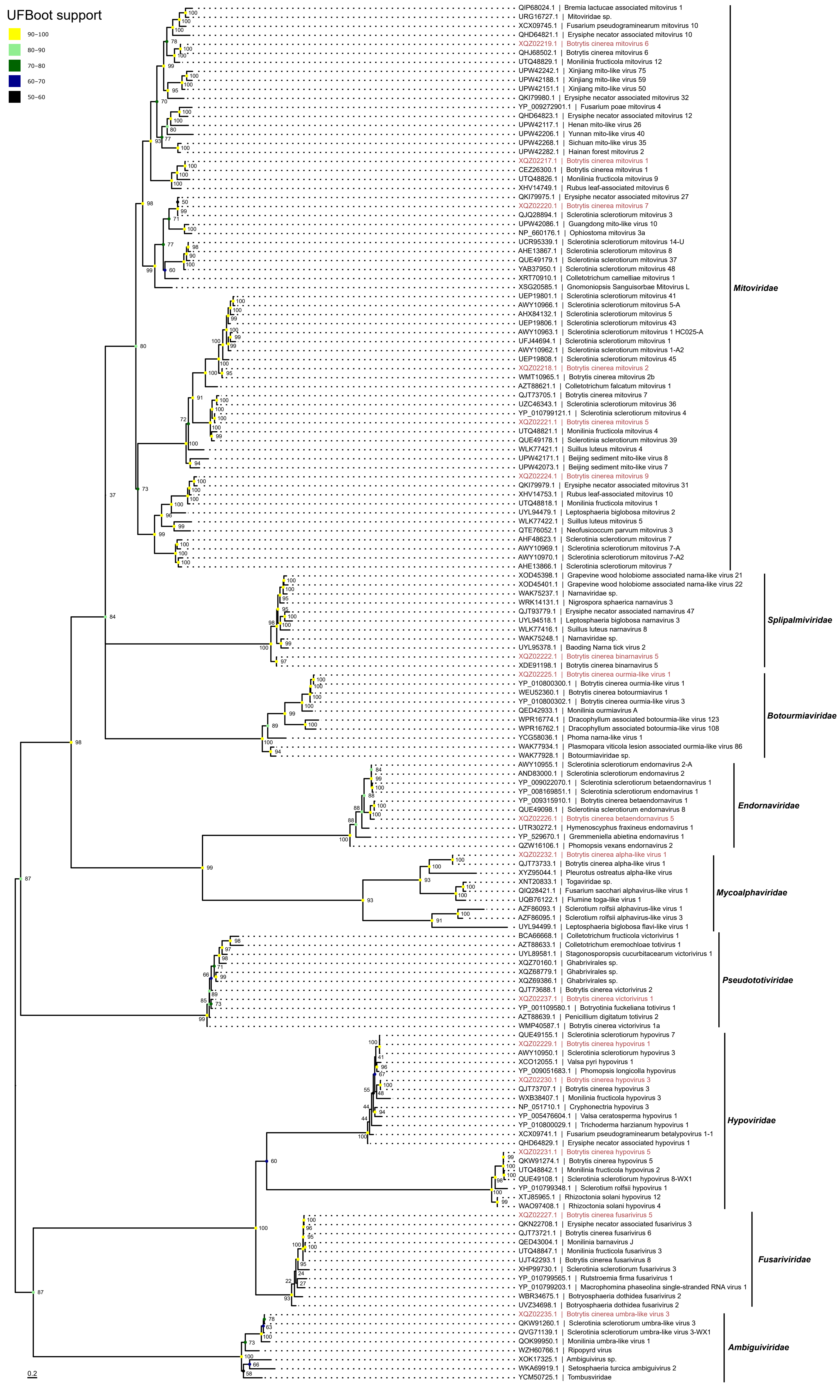

### Supplemental Figure S2

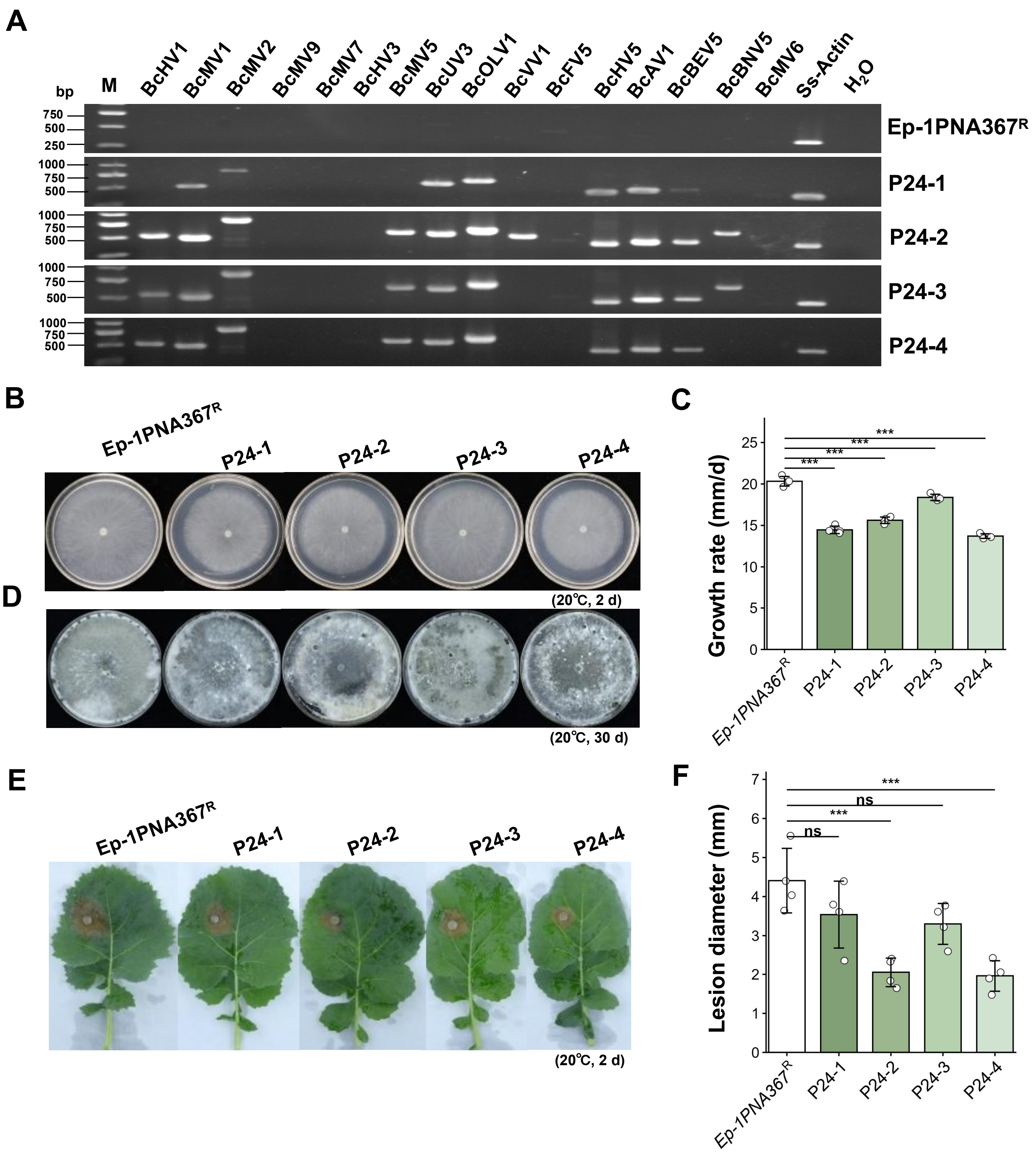
